# Vascular endothelial growth factor-D improves lung vascular integrity during acute lung injury

**DOI:** 10.1101/2024.12.16.628787

**Authors:** Yifan Yuan, Lokesh Sharma, Wenwen Tang, Yongdae Yoon, Shannon Kirk, Micha Sam Brickman Raredon, Farida Ahangari, Johad Khoury, Qian Hong, Yi Luan, Qianying Yuan, Chen Lujia, Yunbo Ke, Konstantin G Birukov, Michael Simons, Dianqing (Dan) Wu, Laura E Niklason, Naftali Kaminski

**Author notes:** These authors contributed equally to the work (Yifan Yuan, Lokesh Sharma, Wenwen Tang, Yongdae Yoon). Corresponding author Corresponding authors: Yifan Yuan.

## Abstract

Disorders in pulmonary vascular integrity are a prominent feature in many lung diseases. Paracrine signaling is highly enriched in the lung and plays a crucial role in regulating vascular homeostasis. However, the specific local cell-cell crosstalk signals that maintain pulmonary microvascular stability in adult animals and humans remain largely unexplored. In this study, we employed single-cell RNA-sequencing (scRNAseq)-based computational pipelines to systematically profile ligand-receptor (L/R) interactions within the lung microvascular niche and identified vascular endothelial growth factor-D (VEGF-D) as a key local factor with previously unrecognized barrier-protective properties in models of acute lung injury. Our scRNAseq data revealed that, under physiological conditions, soluble L/R interactions between mesenchymal cells, in particular alveolar fibroblast, and microvascular endothelial cells are predominantly associated with pathways involved in maintaining vascular integrity as compared to all other cells. Upon treatment with top identified ligands, we found that VEGF-D significantly enhanced endothelial barrier function and conferred protection against inflammatory challenges induced by tumor necrosis factor-α (TNF-α), interleukin-1β (IL-1β), and thrombin. This barrier-protective effect of VEGF-D was significantly attenuated by inhibition of VEGFR2, either through siRNA knockdown or pharmacological blockade using specific VEGFR2 inhibitors. Intravenous administration of recombinant VEGF-D in lipopolysaccharides (LPS)-induced acute lung injury models significantly reduced vascular permeability (7339 ± 2510 (LPS) v.s. 5350 ± 1821 (LPS + VEGF-D), *p* < 0.05), immune cell infiltration (0.791 ± 0.199 x 10^6^ WBC/mL (LPS) v.s. 0.540 ± 0.190 x 10^6^ WBC/mL (LPS + VEGF-D), *p* < 0.01), and the expression of pro-inflammatory markers TNF-α and IL-6 in the lung tissue. This effect was abolished in *VEGFR2^iECKO^* mice, confirming that VEGF-D mediates its effects via VEGFR2-dependent signaling. This study demonstrates an unexpected protective role for VEGF-D in promoting lung endothelial barrier integrity and suggests that paracrine signaling from the alveolar fibroblast niche contributes critically to lung capillary homeostasis.

## 1. Introduction

Vascular homeostasis in the lungs is vital for maintaining the proper function of the pulmonary vascular system, ensuring efficient gas exchange, and preserving the integrity of the alveolar-capillary barrier. The pulmonary vascular monolayer, primarily composed of endothelial cells, forms a critical fluid-tight barrier that facilitates the exchange of oxygen and carbon dioxide in the alveoli (reviewed in (1, 2)). Disruptions in the pulmonary vasculature are linked to various pulmonary diseases. In conditions such as acute lung injury (ALI) and acute respiratory distress syndrome (ARDS), the integrity of the pulmonary endothelium is compromised, leading to increased permeability, edema, and impaired gas exchange (reviewed in (3)). Chronic diseases like chronic obstructive pulmonary disease (COPD) and idiopathic pulmonary fibrosis (IPF) also involve significant alterations in the pulmonary vasculature, contributing to disease progression and severity (reviewed in (2)). In all these diseases, local biochemical and biophysical changes in the vascular microenvironment play a critical role in disease pathogenesis. Understanding these mechanisms is essential for developing therapeutic strategies aimed at preserving or restoring vascular integrity in pulmonary diseases.

Cell-cell crosstalk signaling is central to regulating microvascular stability and maturation, and mesenchymal cells, including smooth muscle cells, pericytes, and fibroblasts, are recognized for their roles in supporting vascular formation and integrity through essential paracrine signals and direct cell-cell interactions (4–7). For example, angiopoietin 1 (Ang-1) and keratinocyte growth factor (KGF), both produced by lung mesenchymal cells, have been shown to enhance barrier function and reduce inflammation in human endothelial cells, preventing the influx of cells and plasma from the bloodstream into the alveoli and airways (reviewed in (8)). In addition, perivascular cells or pericytes reside adjacent to lung capillaries and play a pivotal role in modulating endothelial function and maintaining microvascular stability during lung vascular remodeling through direct cell-cell contact (9). Although the cell-cell interactions in the microvascular niche have been studied mechanistically, primarily during development or in response to discrete injuries, there remains an incomplete understanding of the paracrine signals that regulate the pulmonary microvasculature and maintain it in a homeostatic state in adult animals or humans.

Multi-omics analysis tools provide a comprehensive approach to profiling ligand-receptor (L/R) interactions in multicellular organisms. Previously, we and others developed computational pipelines to analyze single-cell RNA-seq (scRNAseq) dataset and map L/R pairs within and between cell types and assess the significance of these interactions in the animal lung (10–12). In this study, we leveraged these tools to systematically investigate local L/R interactions in the human lung microvascular niche to identify critical targets contributing functional homeostasis and exerting protective impact against injury. The L/R pairs between microvascular endothelial cells (EC) and non-EC lung cells were profiled through *Connectome*, and *NICHES* (10, 11) using previously published single-cell dataset from human lungs, and validated through immunostaining and associated ELISA assays (13). The top ligands were assessed in in vitro cell culture assays to evaluate cell-cell junctions and inflammatory response and in vivo LPS-induced ALI models (Fig. 1A). Our findings revealed that vascular endothelial growth factor-D (VEGF-D), a key soluble factor produced by alveolar fibroblasts, contributes to vascular homeostasis. In both in vitro endothelial cell cultures and in vivo animal models, VEGF-D administration improved barrier integrity and suppressed inflammation, suggesting its potential as a promising pharmacological target for treating lung vascular diseases.

**Figure 1.**
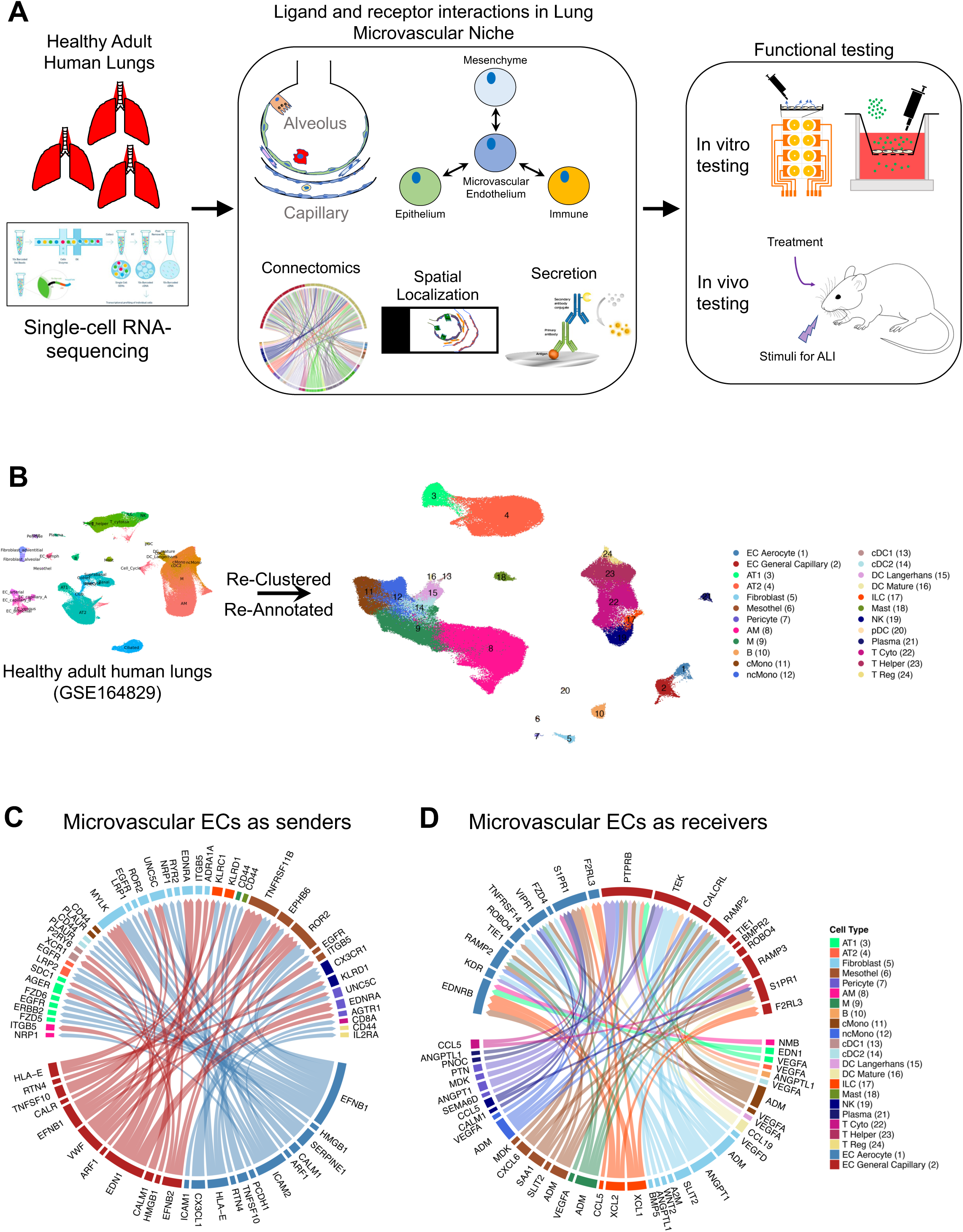
Ligand-receptor interactions between microvascular EC and non-EC lung cells within the human lung microvascular niche. A) Overview of experimental design. First, single-cell RNA-seq data from healthy adult human lung tissue (GSE164829) were analyzed to identify ligand-receptor pairs within the lung microvascular niche. The top identified ligands were validated using immunostaining and ELISA assays. Finally, the biological impact of these ligands was tested in functional in vitro cell culture and in vivo models of acute lung injury (ALI). B) UMAP Plot of the cell populations in the microvascular niche from the human lung. Each dot represents a single cell, and cells are labeled as 1 of 24 discrete cell types. C, D) Circosplot of top 75 edges by edge weight with microvascular ECs as signal senders and receivers.

## 2. Methods

### 2.1. Materials

The following ligands were purchased from vendors: BMP5 (Novus Biologicals, # 615-BMC-020), VEGF-D (Cayman Chemical, #32055), Adrenomedullin (ADM)(Cayman Chemical, #24889), Angiopoietin-1 (Novus Biologicals, #923-AN), TNF-α (R&D Systems, # 10291-TA), Slit homolog 2 (SLIT2)(R&D Systems, # 8616-SL), CXCL6 (R&D Systems, # 333-GC-025/CF), Pleiotrophin (PTN)(R&D Systems, # 252-PL), and Semaphorin 6D (Sema6D)(R&D Systems, #2095-S6).

### 2.1. scRNAseq processing

Single-cell transcriptomic analysis of healthy adult human lung data (GSE164829) was processed through Seurat package (version 3.0 and 4.0). In brief, we applied Variance Stabilizing Transformation (VST) to select the top 2,000 highly variable genes, which were scaled using *ScaleData* function in Seurat for principal component analysis (PCA). Top 30 principal components (PCs) were selected based on the elbow plot generated by the *ElbowPlot* function.

Graph-based clusters were generated by using the *FindNeighbors* and *FindCluster* functions with the selected 30 PCs. All cells across different samples were assigned to two-dimensional Uniform Manifold Approximation and Projection (UMAPs) for visualization. We then applied *FindAllMarkers* function in Seurat to identify differentially expressed genes for each cluster in the object (14). We only chose genes that were expressed in at least 10% of cells in one of these clusters to ensure the quality of genes. These marker lists were used for re-annotation based on top defining genes of canonical cell types in the literature (13, 15, 16). The data was then subsetted using the *subset* function in the Seurat package to include only the cell populations present in the microvascular niche, removing cell populations from large airways, large vessels, lymphatics, and multiplets.

### 2.2. ScRNAseq cell-cell communication Analysis

To study the cell-cell communication within the human lung microvascular niche, data was mapped to the NicheNet ligand-receptor interaction database (11) using the R software Connectome (v1.0.0) (https://msraredon.github.io/Connectome/)(10, 17)(Supplemental Table 1). Selected ligand-receptor pair were visualized as *Circosplots* after data filtration based on the criteria used previously (13): Ligands being significantly expressed in Microvascular EC, such as EC Aerocyte Capillary and EC General Capillary and receptors significantly expressed in non-EC populations or vice versa (using Bonferroni adjusted p-value threshold of p_adjust < 1e^-5^). Ligands and receptors were expressed in at least 20% of cells in their respective cell types, with mean scaled expression values > 0 were included in the analysis. We removed integrin receptors, extracellular matrix (ECM) ligands, cell-cell junction, and adhesion signaling molecules in our analysis due to their insoluble nature or possible contribution from a non-cellular origin. The top 75 interactions ranked by scaled weight were shown using *Circosplots* function in Figure 2. Analysis of ligand-receptor signaling between mesenchymal cells and microvascular endothelial cells was performed using NICHES v1.0.0 as previously described (18).

**Figure 2.**
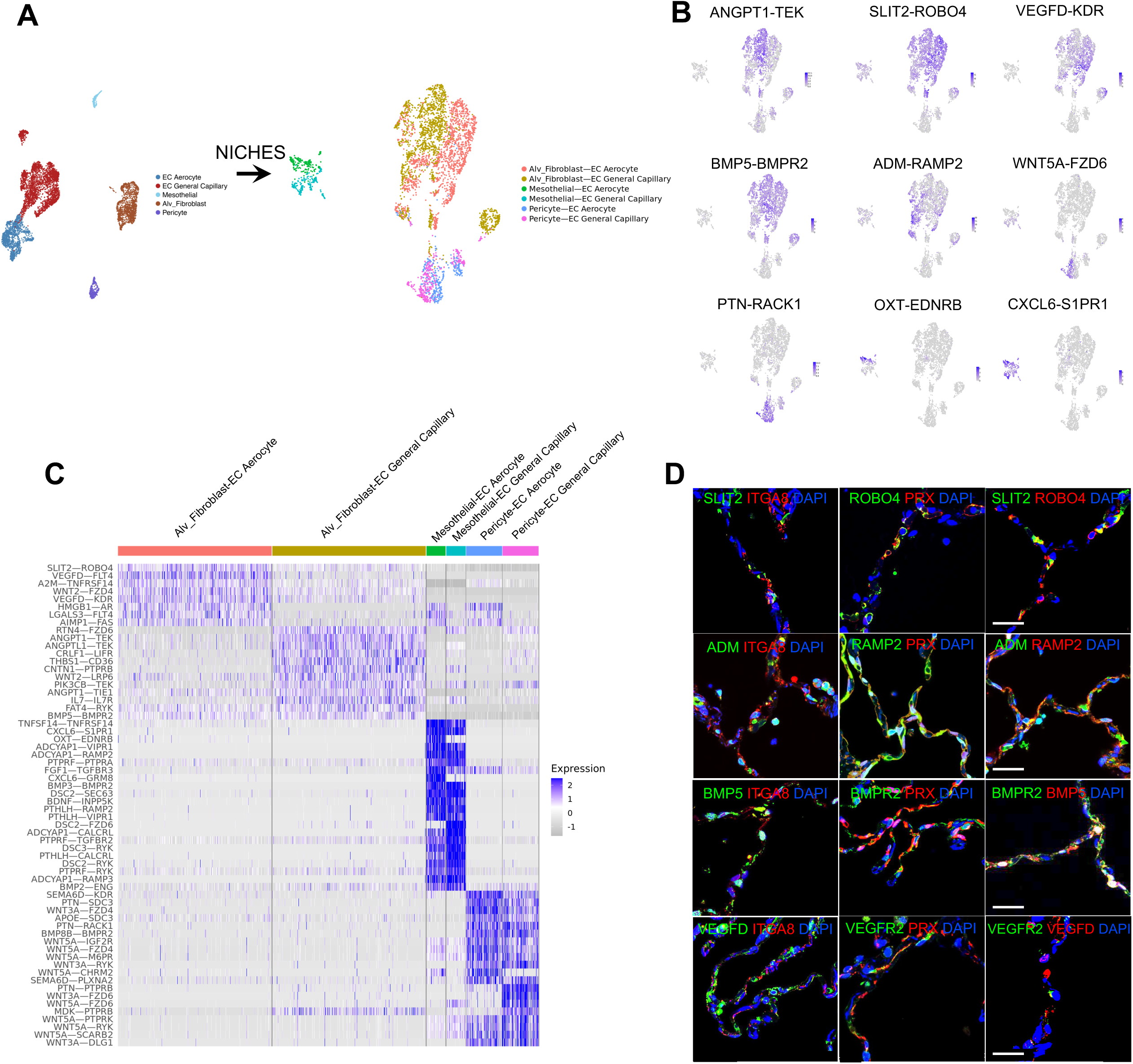
Distinct L/R pairs between mesenchymal cell and microvascular EC within the human lung microvascular niche. A) NICHES analysis of AFs, mesothelial cells, pericytes, gCap, and aCap in human microvascular niche yields a quantitative cell-cell signaling atlas of AF-aCap, AF-gCap, Mesothelial-aCap, Mesothelial-gCap, Pericyte-aCap, and Pericyte-gCap visualized by low-dimensional UMAP embedding. B) FeaturePlot of top differentially expressed L/R pairs *ANGPT1*-*TEK*, *SLIT2*-*ROBO4*, *VEGFD*-*KDR*, *BMP5-BMPR2*, *ADM*-*RAMP2*, *WNT5A*-*FZD6*, *PTN*-*RACK1*, *OXT*-*EDNRB*, and *CXCL6*-*S1PR1* from NICHES analysis. C) Top 10 differentially expressed L/R pairs between different cell-cell interaction signals processed through NICHES and plotted in DoHeatmap. D) Immunostaining of representative L/R pairs from AF to microvascular endothelial cells, including *SLIT2*-*ROBO4*, *ADM*-*RAMP2*, *BMP5*-*BMPR2*, and *VEGFD*-*VEGFR2* in human lung section. *ITGA8* is a marker for AF; *PRX* is a marker for microvascular endothelial cells. Scale bar 20 μm.

### 2.3. Immunofluorescent Staining

Immunofluorescent staining was performed as previously described (13, 19, 20). Native FFPE (formalin fixed paraffin embedded) human lung sections were de-parafinized through soaking in antigen retrieval solution (0.05% Tween-20, 1mM EDTA, and 10 mM Tris (pH = 9)) at 75 °C for 20 minutes, followed by blocking (10 % FBS and 0.2% Triton X-100, RT, 20 minutes). Tissue samples were then stained with antibodies: SLIT2 (Thermo Fisher Scientific, # PA5-31133), ROBO4 (R&D Systems, # AF2366), ADM (Bioss, # BS-0007R), RAMP2 (Santa Cruz Biotechnology, # sc-365240), BMP5 (Thermo Fisher Scientific, # PA5-78878), BMPR2 (Thermo Fisher Scientific, # MA5-15827), VEGFD (Thermo Fisher Scientific, # PA5-13300), VEGFR2 (Santa Cruz Biotechnology, # sc-6251), ITGA8 (Novus Biologicals, # AF4076), PRX (Novus Biologicals, # NBP1-89598) overnight at 4 °C followed by staining with secondary antibodies (RT, 1 hour) and DAPI (RT, 5 min).

To detect cytoskeletal protein and tight junctions, human lung microvascular endothelial cells (HMVECs)(Lonza, # CC-2527) were seeded into fibronectin-coated 12-well plate at a density of 1 × 10^4^ cells/cm^2^. Once reaching confluence, adherent cells were fixed with 2% paraformaldehyde (containing Ca^2+^/Mg^2+^) (RT, 15 min), permeabilized with 0.1% Triton-X (RT, 3 min), and blocked with 5% FBS 1XPBS for 1 hour at RT. Cells were then stained with VE-Cadherin (Abcam, # ab33168) overnight at 4 °C. This was followed by a 1-hour incubation with a secondary antibody conjugated with Alexa Fluor 488, followed by staining with rhodamine phalloidin (20 min, ThermoFisher Scientific, # R415) and DAPI (5 min). All samples were washed, mounted, and assessed with confocal microscopy (Leica SP5).

### 2.4. Quantitative Real-time Reverse Transcription-PCR (qRT-PCR)

Methods for qRT-PCR were performed as previously described (20). Briefly, total mRNA was extracted from cells using an RNA isolation kit (Qiagen) and cDNA was synthesized (Bio-Rad). The mRNA levels of human genes including *ICAM1*, *VCAM1*, *IL6*, *CXCL5*, *TNF*, *IL8*, and *GAPDH* were analyzed using SYBR green primers (IDT Technologies) and a Real-Time PCR system (Bio-Rad). The sequences of each primer are listed in Table 1.

**Table 1.**
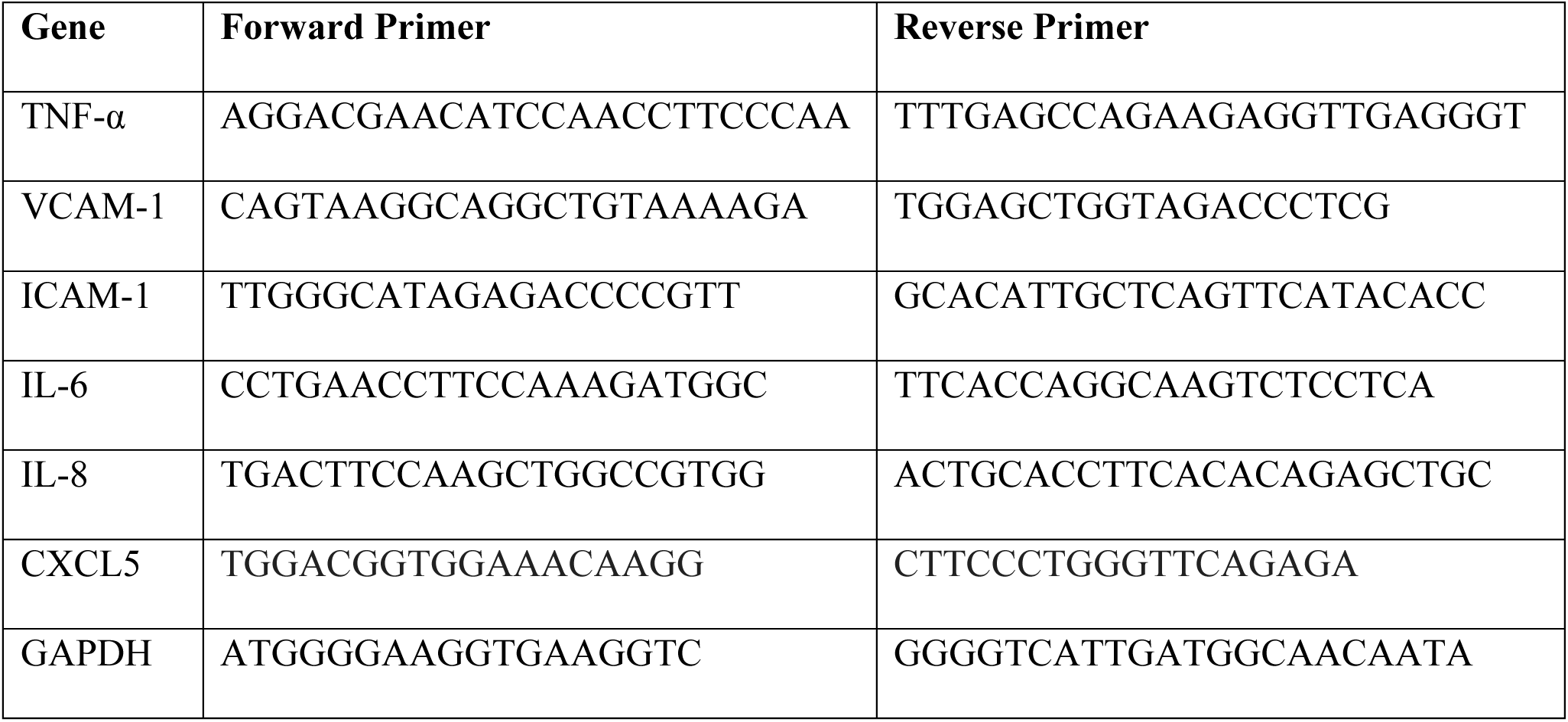
List of primer sequences used for RT-PCR.

### 2.5. ELISA assay

To determine the secretion level of the top ligands identified from scRNAseq data, ex vivo rat lung culture were used as a surrogate model. Healthy adult rat lungs (8 – 10 weeks) were extracted and minced into small pieces (<1mm^2^) followed by culturing in DMEM medium supplemented with 2% FBS for a total of 20 hours. Supernatant were taken at 1 hour, 6 hours, and 20 hours from the ex vivo lung pieces culture. The protein levels of SLIT2, ADM, BMP5, and VEGFD in samples harvested from different timepoints were determined using corresponding ELISA kits: SLIT2 (# NB030384), ADM (# NBP2-78738), BMP5 (# NBP2-69996), VEGFD (# NBP2-78890), all from Novus Biologicals. The OD values in all samples were read in Spectra MAX® iD5 (Molecular Deices).

To quantify the soluble ICAM-1 protein level from culture medium, we used Human ICAM-1 ELISA kit (#DY720) with Clear Microplate (#DY990) from R&D systems (Minneapolis, MN, USA). All ELSIA assays were conducted according to the manufacturer’s instructions. The measurement was quantified using Spectra MAX® iD5 (Molecular Deices)

### 2.6. Electrical cell-substrate impedance sensing

Electrical cell-substrate impedance sensing (ECIS) assay was performed as previously described (20). In brief, the ECIS plates were first functionalized with 10mM L-cysteine followed by coating with fibronectin (1 µg/cm^2^). HLMVECs (Lonza, # CC-2527), human pulmonary vein endothelial cells (HPVECs) (Cell Biologics, # H-6060), or human pulmonary artery endothelial cells (HPAECs) (Lonza, # CC02530) were seeded into the ECIS plate at a concentration of 50,000 cells/cm^2^. Ligands were added 2 – 3 days after seeding at which resistances reached plateau. The transendothelial electrical resistance (TEER) was measured and calculated by ECIS system at real time.

### 2.7. Transwell Assay

Transwell assay was performed as previously described (20). In brief, HLMVECs were seeded into fibronectin-coated transwell inserts (polyester, 6.5 mm in diameter, pore size 0.4 µm) at a density of 50,000 cells/cm^2^. After 2 – 3 days of culture, individual ligand was added into the upper well of the inserts for 30 minutes followed by replacing the medium with 150 µL of FITC-Dextran (1 mg/mL, Sigma-Aldrich, 20 kDa, #FD20S) containing medium into the inserts. After another 30 minutes of incubation, samples were collected from the lower compartment of the transwell insert. The concentration of FITC-Dextran in different samples were determined by spectrophotometer at a wavelength of 488 nm. To evaluate the response to pro-inflammatory cytokines, cells were challenged with TNF-α (1 ng/mL, 5 hours), IL-1β (5 ng/mL, 5 hours), or thrombin (10 U/mL, 20 minutes) prior to treating with VEGF-D. The FITC-dextran was added after 30 minutes treatment with VEGF-D.

### 2.8. Western Blotting

The cells were lysed by 2x Laemmli Sample Buffer (Bio-rad, # 1610737), boiled for 5 min, subjected to SDS-polyacrylamide gel electrophoresis, and transferred to polyvinylidene difluoride membrane (PVDF, Millipore). The membrane was blocked with 5% skim milk or bovine serum albumin in Tris-HCl-buffered saline containing 0.05% Tween 20 (TBST) for 30 min and incubated with primary antibodies such as VCAM-1 (#13662), IκBα (#4814), NF-κB p65 (#4764), and Thr18/Ser19 Phospho-Myosin Light Chain 2 (#3674) from Cell Signaling Technology (Danvers); ICAM-1 (Santa Cruz Biotechnologies, # sc-8439,); Beta-actin (ThermoFisher Scientific, #MA5-15739) for overnight at 4℃. After washing with TBST, the membrane was incubated with horseradish peroxidase-conjugated secondary antibodies (Cell signaling Technology, #7074 and #7076) for 1 hour. After another washing with TBST, the bands were visualized with SuperSignal^TM^ West Pico PLUS Chemiluminescent Substrate (# 34580) or SuperSignalTM West Atto Ultimate Sensitivity Substrate (# A38556) from ThermoFisher Scientific, and the images were generated using a ChemiDoc XRS+ System (Bio-Rad).

### 2.9. siRNA transfection

HLMVECs were seeded at 10,000 cells/cm^2^ into 6-well plate and cultured overnight to reach 50 – 60% confluence. This was followed by transfection with siRNA: VEGFR2 siRNA (Santa Cruz Biotechnologies, # sc-29318) or control siRNA (Santa Cruz Biotechnologies, # sc-37007) as per manufacturer’s instructions (Lipofectamine RNAiMAX, Thermo Fisher Scientific, # 13778075). The media was then changed with complete cell culture medium (VascuLife® VEGF Endothelial Medium (LifeLine, # LL-0003) supplemented with 12.5% FBS and 1.5% Antibiotics/Antimycotics) after 6 hours of transfection.

### 2.10. LPS-induced acute lung injury animal model

The LPS-induced acute lung injury model was established as previously described (21). In brief, we used female C57BL/6 mice, aged 7 – 8 weeks, obtained from Envigo. All procedures were conducted following the guidelines of the Institutional Animal Care and Use Committees of Yale School of Medicine. Mice were anesthetized with combination of ketamine and xylazine.

LPS at 0.25 mg/kg was administered intra-tracheally to mice to induce acute lung injury. Recombinant mouse VEGF-D protein at 225 µg/kg (Bio-techne, #469-VD-025/CF) or saline control was administered retro-orbitally about 30 minutes after LPS injection. After another 18 hours, mice under ketamine/xylazine anesthesia were first subject to BAL (Bronchoalveolar lavage) collection and then were euthanized by tissue (blood) and vital organ (lungs) harvesting for later analysis.

### 2.11. Bronchoalveolar lavage fluids (BAL)

BAL samples were prepared as described in previous study (21). Briefly, after euthanizing the mice, trachea was exposed and dissected, and a 20-gauge (×1¼ in) catheter was inserted. The lungs were gently lavaged twice with 1 ml of sterile non-pyrogenic phosphate-buffered saline solution (PBS, pH 7.4). BAL samples from each mouse were centrifuged at 5,000 rpm for 10 minutes at 4°C. The cell pellets were resuspended in 300 µL of 1X PBS, then counted using an automated cell counter or hemacytometer. The supernatants were collected and stored at −80°C for subsequent analysis of total protein content using a bicinchoninic acid (BCA) assay (Bio-rad, # 5000001). Following the lavage, the left lung lobes were harvested for histology, and the right and accessory lobes were stored at −80°C for subsequent analysis.

### 2.12. Mice

*VEGFR2^iECKO^* was generated by crossing *VEGFR2^e3loxP/e3loxP^* (stock number 018977, Jackson Laboratory) with C57BL/6-Tg (Cdh5-cre/ERT2)1Rha originally from Dr. Ralf Adams lab (22, 23). The *VEGFR2^iECKO^*mice were a general gift from Dr. Michael Simons lab at Yale School of Medicine. All animal husbandry and experiments were approved by the Institutional Animal Care and Use Committees at Yale University in accordance with US National Institutes of Health guidelines, and in accordance with U.S. Government Principles for the Utilization and Care of Animals Used in Research, Teaching and Testing. Mice were maintained on a C57BL/6 background and genotyped to determine their control versus appropriate lineage tracing genotypes and allocated accordingly at weaning. Both sexes were used for this study.

### 2.13. Statistical Analysis

Comparisons between groups were performed with Student’s *t-*test or, for multi-group comparisons, by analysis of variance (ANOVA) followed by a post-hoc test. All analysis was performed with GraphPad Prism 10.0; values shown graphically are mean ± standard error.

## 3. Results

### 3.1. Profiling Ligand-Receptor Interactions between Microvascular EC with non-EC Lung Cells in Human Lung

To specifically analyze cell populations within the pulmonary microvascular niche, we excluded cells from large vessels, such as pulmonary arterial endothelial cells, pulmonary venous endothelial cells, systemic venous endothelial cells, vascular smooth muscle cells, and cells from the large and small airways, including basal, suprabasal, goblet, club, ciliated, ionocyte, PNEC, airway smooth muscle cells, and lymphatic cells. We then reanalyzed and reannotated our previously published scRNAseq dataset (13) (Fig. 1B), which comprises samples from 68 healthy adult individuals, including 29 females and 39 males, with a median age of 50 (range: 32–61 years). The final dataset includes a total of 221,171 cells, categorized into 24 distinct cell populations within the pulmonary microvascular niche, each expressing previously validated canonical markers (Supplemental Fig. 1). The sequencing data has been previously made publicly available on Gene Expression Omnibus under the accession number (GSE164829).

To profile L/R interaction signals between microvascular endothelial cells (EC) and non-EC lung cell populations, we applied *Connectome* to map the average gene expression per cell type to known L/R interactions cataloged in the NicheNet database (10, 11)(Supplemental Table 1). The analysis focuses on the interaction between pulmonary microvascular ECs and non-EC cell types and only the secreted ligands are included (Figs. 1C and D). Using microvascular EC subtypes, specifically EC Aerocyte (aCap) and EC General Capillary (gCap), as signal senders (Ligands) and the adjacent cell types (alveolar fibroblasts, AT1, AT2, pericytes, plasma cells, etc.) in the microvascular niche as signal receivers (Receptors), we identified several abundant and canonical L/R pairs involved in immune response, including *HLA-E* and *TNFSF10*, expressed by both gCap and aCap ECs and interacted with *KLRC1*, *KLRD1*, and *TNFRSF11B*, which were expressed by innate lymphoid cells (ILC), natural killer (NK), and mesothelial cells, respectively ((24) and reviewed in (25)). Specifically, gCap ECs produced *EDN1*, which binded with *EDNRA* expressed by pericytes and alveolar fibroblasts (AFs), acting as a vasomotor (13, 26). Additionally, aCap ECs uniquely contributed to blood clot regulation by secreting *SERPINE1*, which modulated urokinase plasminogen activator receptor (PLUAR) expressed by cDC2 and classical monocytes (cMono) cells, alongside exhibiting low levels of von Willebrand factor (VWF) (Fig. 1C).

Using aCap and gCap as signal receivers (receptors) and adjacent pulmonary cell populations as signal senders (ligands), we identified a group of abundant L/R interactions critical for vascular junction stability and anti-inflammatory responses. Notably, these interactions such as *SLIT2-ROBO4*, *ANGPT1-TEK*, and *ADM-RAMP2* were enriched between mesenchymal cells, such as AFs, pericytes, and mesothelial cells, and aCap and gCap endothelial cells, and were known to contribute to the regulation of vascular integrity (27–30). Additionally, pro-angiogenic signals like pleiotrophin (PTN)-*PTPRB* and midkine (MDK)-*PTPRB* (reviewed in (31)) were highly expressed between gCap and pericytes, as well as mesothelial cells, suggesting an active role of these signaling axes in neovascularization in the lung microvasculature, particularly within gCap-enriched niche. Interestingly, aCap exhibited distinct and high expression of the vasodilatory receptors *EDNRB* and *VIPR1*, indicating their role as principal signal receivers for vasodilatory ligands such as *EDN1*, secreted by alveolar type 1 (AT1) cells, and adrenomedullin (ADM), produced by AFs, mesothelial cells, and interstitial macrophages (Fig. 1D). Additionally, an enriched expression of *VEGFA-KDR*, and *VEGFA-PTPRB* pairs between various non-EC lung cells to microvascular endothelial cells was found in the microvascular niche, confirming the important role of VEGFA signaling in contributing vascular homeostasis (reviewed in (32)).

### 3.2. Alveolar fibroblasts express ligands to regulate vascular integrity

To elucidate distinct L/R interactions between mesenchymal cells and microvascular endothelium, we applied *NICHES* (18) to systematically analyze differentially expressed L/R pairs, focusing on signals from AFs, mesothelial cells, and pericytes, targeting the endothelial subtypes aCap and gCap (Fig. 2A). Our analysis revealed that AFs express high level of L/R pairs such as *SLIT2-ROBO4* (AFs to aCap), *ANGPT1-TEK* (AFs to gCap), and *ADM-RAMP2* (AFs to aCap and gCap), which are critical in regulating vascular junction integrity, compared to mesothelial cells and pericytes (Figs. 2B and C). In contrast, pericytes predominantly expressed ligands associated with neovascularization, including *PTN*, *MDK*, and *SEMA6D*, as well as WNT signaling (*WNT5A*, *WNT3A*), which interacted with receptors *FZD6* in gCap and *FZD4* in aCap to regulate processes such as cell polarization, mechanotransduction, and alveolar homeostasis (Figs. 2B and C)(33–35). Mesothelial cells were found to uniquely produce vasodilatory signals, such as *ADCYAP1-CALCRL*, *ADCYAP1-RAMP2*, *CXCL6-S1PR1*, *OXT-EDNRB*, and *PTHLH-VIPR1*, interacting with both gCap and aCap to modulate vascular tone (Figs. 2B and C)(36). These findings underscore the distinct and specialized roles of mesenchymal cell subtypes in modulating endothelial behaviors and suggests paracrine signals from AFs may exert function to improve vascular stability.

To further confirm the L/R interactions at the protein level, spatial localization of the top identified pairs was determined in native human distal lung samples. Canonical markers, such as *ITGA8* and *PRX*, were used to label AFs and microvascular endothelial cells, respectively (13). Immunostaining images revealed that top L/R pairs, including SLIT2-ROBO4, VEGFD-VEGFR2, BMP5-BMPR2, and ADM-RAMP2, were either co-localized or adjacent, suggesting a strong possibility of crosstalk (Fig. 2C). To further determine the secretion level of the identified ligands, we measured the protein level in the supernatant from an ex vivo rat lung culture as a surrogate model (Supplemental Fig. 2). The lungs were extracted from adult rats, cut into small pieces (<1mm³) with surgical scissors, and cultured in serum-reduced medium for 20 hours. A small volume of supernatant was collected at 1, 6, and 20 hours from the same well. The levels of SLIT2, ADM, BMP5, and VEGFD in the supernatant reached biologically meaningful levels as early as one hour (Supplemental Fig. 2). The combination of computational analysis with biochemical assays allowed to identify a group of high-fidelity L/R pairs with validated localization and secretion, and with high likelihood of interaction with pulmonary microvascular endothelium.

### 3.3. VEGF-D improves vascular integrity in human lung microvascular endothelial cells cultured in vitro

To evaluate the impact of putative ligands produced by AFs on endothelial functions, we used electrical cell impendence sensing (ECIS) and transwell assays to measure permeability.

Human lung microvascular endothelial cells (HLMVECs) were cultured for two days in an ECIS plate to reach confluence, after which they were treated with top ligands from AFs, including ADM, Ang-1, SLIT2, VEGF-D, BMP5, and top ligands from mesothelial cells and pericytes including CXCL6, semaphorin 6D (SEMA6D), and pleiotrophin (PTN). Consistent with previous findings (37–39), ADM and Ang-1 significantly enhanced transendothelial electrical resistance in the endothelial cell monolayer, indicating an improvement in barrier function. In contrast, other ligands, including SLIT2, BMP5, CXCL6, SEMA6D, and PTN, had either modest or negligible effects on endothelial barrier integrity (Fig. 3A).

**Figure 3.**
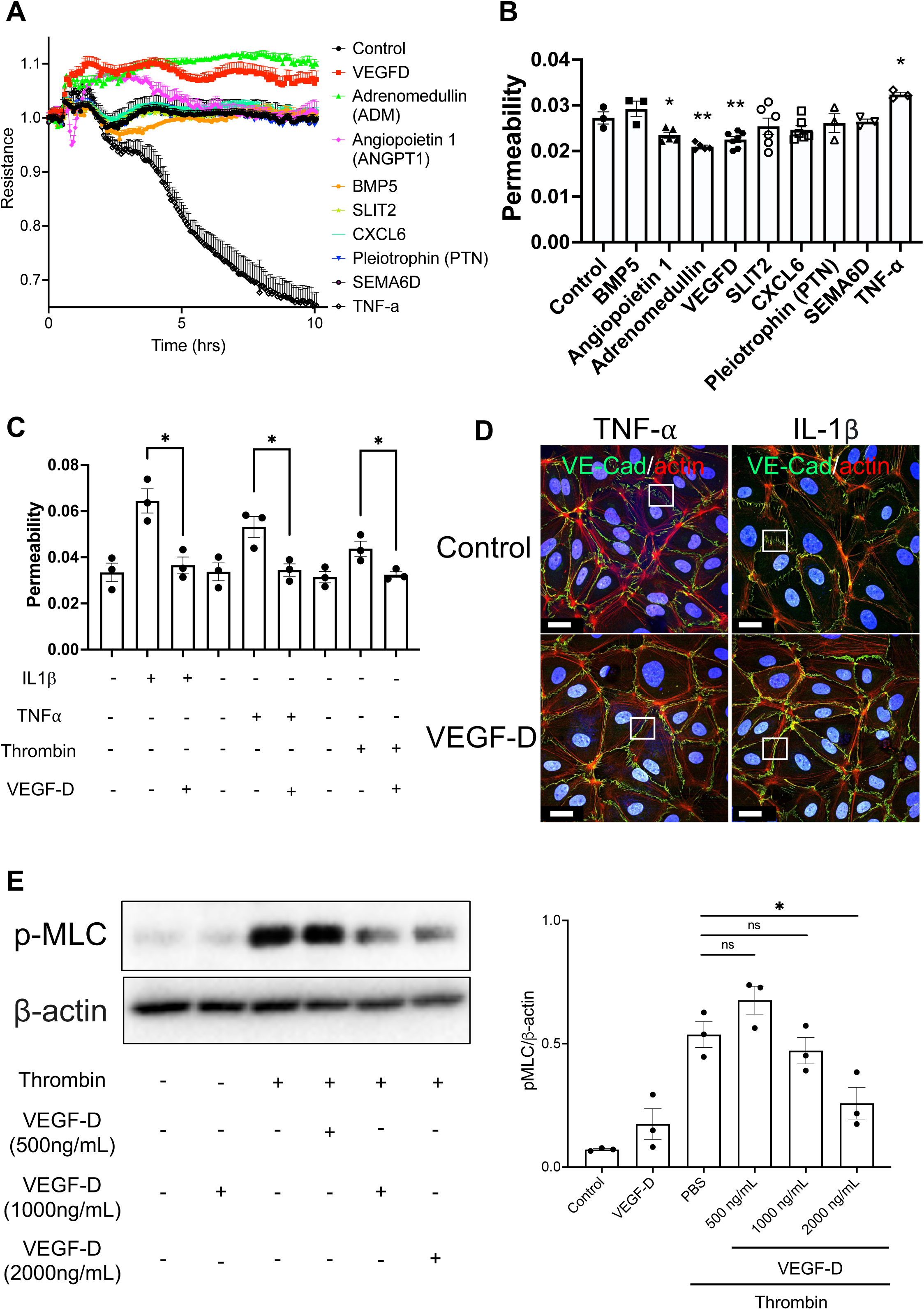
VEGF-D increases the barrier function of HLMVECs. A, B) The electrical resistance and permeability in HLMVECs after treatment with VEGF-D (1 μg/mL), adrenomedullin (ADM)(100 ng/mL), angiopoietin 1 (ANGPT1)(2 μg/mL), BMP-5 (1 μg/mL), SLIT2 (1 μg/mL), CXCL6 (1 μg/mL), Pleiotrophin (PTN)(1 μg/mL), SEMA6D (1 μg/mL), and TNF-α (1 ng/mL) were measured through electrical cell impendence sensing (ECIS) assay, and transwell assay, respectively. C) HLMVECs cultured on transwell treated with VEGF-D, followed by challenging with IL1β, TNF-α, or thrombin. The permeability was determined using a 20kDa FITC-dextran probe. D) Immunofluorescent images of F actin (red), and VE-Cadherin (green) in HLMVECs challenged with TNF-α or IL1β with or without VEGF-D. White boxes highlight VE-Cadherin junctions in all conditions. Scale bar 20 μm. E) Protein expression of phosphor-MLC in HLMVECs challenged with thrombin with different concentrations of VEGF-D was assessed through a western blot. Data presented as mean ± SEM. * and ** indicate p<0.05 and p<0.01, respectively.

Interestingly, VEGF-D, previously known as a potent activator of endothelial cells through its role in promoting angiogenesis and lymphangiogenesis (40, 41), was found to significantly increase electrical resistance in HLMVECs (Fig. 3A), as well as in human pulmonary vein and arterial endothelial cells (Supplemental Figs. 3A and B). Consistently, in the transwell assay, VEGF-D-treated cells demonstrated significantly lower permeability (FITC-dextran flux: 0.0230 ± 0.00150 (VEGF-D) vs. 0.0273 ± 0.00232 (control), *p* < 0.01) (Fig. 3B), suggesting that VEGF-D may play a novel role in enhancing endothelial barrier integrity and maturation under standard in vitro conditions.

To assess the protective effects of VEGF-D on barrier function during endothelial injury, HLMVECs were pre-treated in transwell inserts with VEGF-D before challenged with barrier-disruptive stimuli, including IL1β, TNF-α, or thrombin. VEGF-D treatment significantly reduced endothelial permeability following challenges with IL1β (0.0644 ± 0.00909 (IL1β) vs. 0.0367 ± 0.00608 (IL1β + VEGF-D), *p* < 0.05), TNF-α (0.0531 ± 0.00795 (TNF-α) vs. 0.0348 ± 0.00467 (TNF-α + VEGF-D), *p* < 0.05), and thrombin (0.0438 ± 0.00565 (thrombin) vs. 0.0324 ± 0.00229 (thrombin + VEGF-D), *p* < 0.05) (Fig. 3C).

The reorganization of stress fibers and cortical actin is closely linked to cell-cell junction regulation (reviewed in (42)). Immunostaining of HLMVECs revealed that the distribution of cortical F-actin and junction protein VE-Cadherin at the cell membrane transitioned from a discontinuous and punctuated pattern during TNF-α or IL1β challenges to a more continuous pattern at the cell periphery following VEGF-D treatment (Fig. 3D). This suggests that VEGF-D may enhance vascular barrier integrity through the reorganization of the actin cytoskeleton.

Furthermore, we observed that myosin light chain phosphorylation (p-MLC), a driver of actomyosin contractility and stress fiber formation, which disrupts the endothelial barrier (43), was significantly increased after thrombin challenge but was downregulated following VEGF-D treatment (0.537 ± 0.0902 (thrombin) vs. 0.259 ± 0.111 (thrombin + VEGF-D), *p* < 0.05) (Fig. 3E), confirming the involvement of actin cytoskeleton-mediated mechanisms. Notably, there were no changes in the gene expression levels of key cell-cell junction proteins, including VE-Cadherin, ZO1, Occludin, and Claudin-5, following VEGF-D treatment (Supplemental Fig. 3C). These findings indicate that VEGF-D exerts a protective effect on HLMVECs against barrier disruption, primarily through mechanisms dependent on the actin cytoskeleton.

### 3.4. VEGF-D resolves vascular inflammation in human lung microvascular endothelial cells

To investigate the effect of VEGF-D on the endothelial inflammatory response induced by cytokines, HLMVECs were pre-treated with VEGF-D before being exposed to TNF-α for 5.5 hours. Following treatment, the mRNA levels of inflammatory cytokines and chemokines, including *CXCL5*, *TNF*, as well as endothelial cell adhesion molecule *ICAM1*, were significantly reduced (Fig. 4A). The NF-κB pathway is a classic signaling cascade that is involved in endothelial inflammation (reviewed in (44)). Western blot analysis showed a notable increase in the levels of the inhibitory NF-κB subunit IκBα after VEGF-D treatment (Fig. 4B), indicating that VEGF-D plays a protective role by preventing the degradation of IκBα and the subsequent canonical activation of the NF-κB pathway induced by TNF-α. Moreover, VEGF-D treatment led to a dose-dependent reduction in the protein levels of total ICAM-1 in cell lysates, as well as soluble ICAM-1 measured by ELISA (Figs. 4B and C). These findings demonstrate that VEGF-D treatment effectively suppresses the inflammatory response triggered by TNF-α in an in vitro cell culture system.

**Figure 4.**
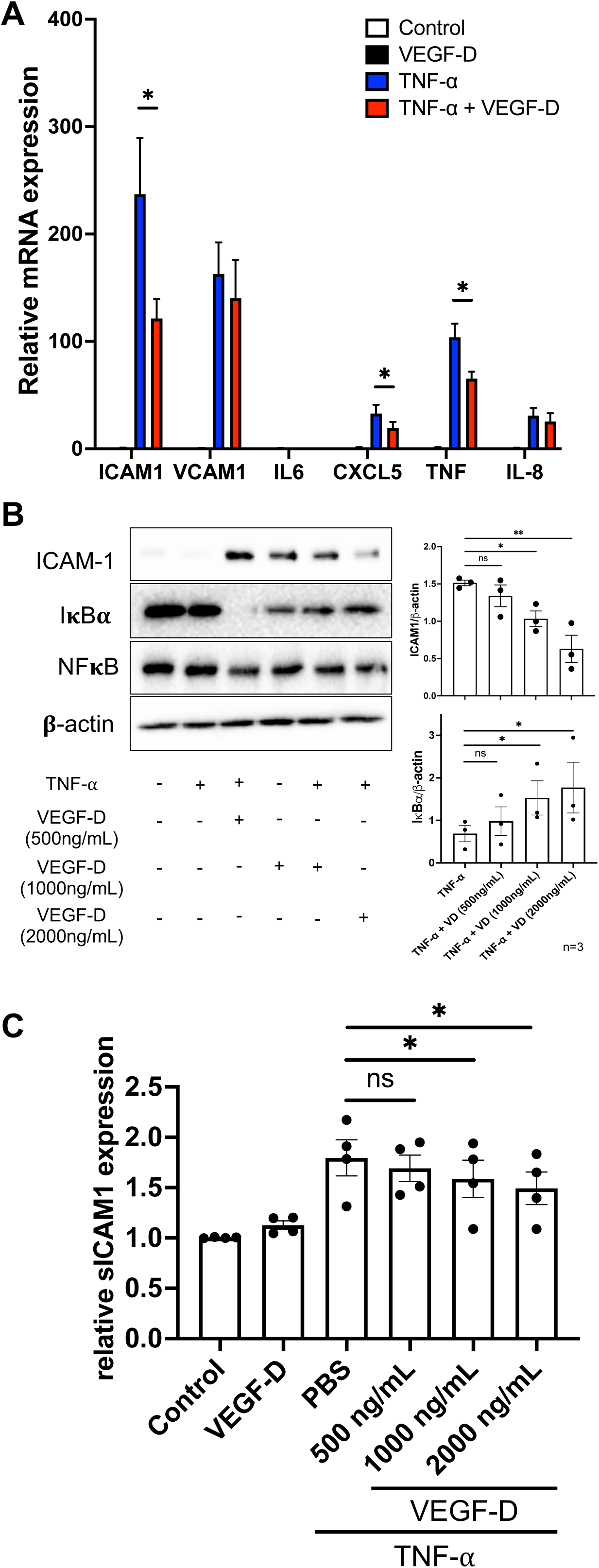
VEGF-D suppresses pro-inflammatory response from TNF-α challenge. A) HLMVECs were treated with VEGF-D for 45 minutes followed by TNF-α (1 ng/mL) challenge. Samples were collected after 5.5 hours of treatment with TNF-α. The gene expression levels of *ICAM1*, *VCAM1*, *IL6*, *CXCL5*, *TNF*, and *IL8* in all conditions was measured through qRT-PCR. Samples with no TNF-α challenge were used as control. B) HLMVECs were treated with different concentrations (500 ng/mL, 1 μg/mL, and 2 μg/mL) of VEGF-D for 45 minutes, followed by a TNF-α challenge. Protein expression of ICAM-1, IκBα, and NFκB was determined by western blot. The densitometry analysis was performed through ImageJ. C) HLMVECs were pre-treated with different concentrations of VEGF-D followed by a TNF-α challenge. The supernatant of different conditions was collected to detect soluble ICAM1 levels through ELISA assay. Data presented as mean ± SEM. * and ** indicate p<0.05 and p<0.01, respectively.

### 3.5. VEGF-D alleviates vascular leakage during acute lung injury

To investigate the role of VEGF-D in lung vascular injury, we first examined the expression levels of VEGFD in healthy versus ALI animal models. Previous studies have shown that administering LPS to the lungs induced inflammation, compromised vascular permeability, increased neutrophil infiltration, and led to pulmonary edema, mimicking ARDS (reviewed in (45)). In our LPS-induced lung injury models, we observed a significant decrease in VEGFD levels in BAL fluid compared to saline controls after 18 hours of treatment with LPS (Fig. 5A). scRNAseq data reveals that *VEGFD* is primarily expressed by AFs in both human and mouse lungs, evidenced by its co-expression with canonical markers of AF in both species, including *ITGA8*, *GPC3*, and *FGFR4* for human, and *Col13a1*, *Tcf21*, *Itga8*, *Npnt*, and *Wnt2* for mouse (46) (Supplemental Figs. 4A and B). We then evaluated the *VEGFD* expression level in the AF population using the scRNAseq data in the bleomycin-induced acute lung injury animal model from previously published dataset (47). Our analysis revealed that the transcriptomic level of *Vegfd* in tdtamato+ AF population in *Scube2-creER Rosa26-tdTomato* mice (47) was gradually decreased overtime between day 0 to day 21 after bleomycin injection (Supplemental Fig. 5A). Additionally, the expression of *Vegfd*, *Angpt1*, and *Slit2* were dramatically decreased with the phenotypic shifting from normal to pro-inflammatory (*Ccl2, Saa3, Cxcl2* (47)) and to fibrotic fate (*Cthrc1, Col1a1, Postn* (47, 48)) (Supplemental Fig. 5B) in AF population. These findings demonstrate that VEGFD expression is reduced during animal models of acute lung injury and suggest its potential role in maintaining tissue homeostasis.

**Figure 5.**
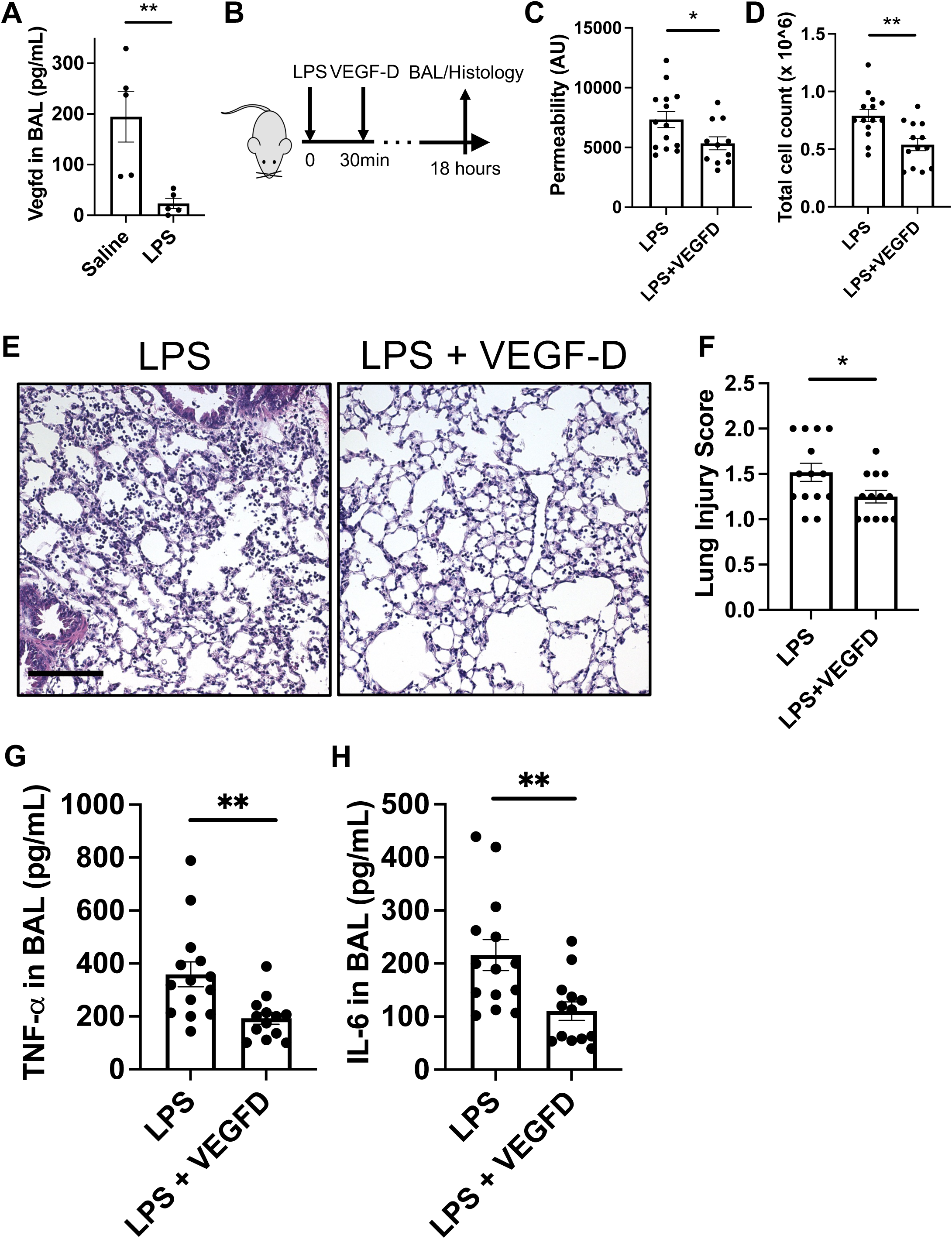
VEGF-D reduces vascular leakage and resolves inflammation in LPS-induced acute lung injury. A) C57Bl/6 mice were injected with LPS for 18 hours, followed by a collection of bronchioalveolar lavage (BAL) fluid. The expression level of VEGF-D was measured in the BAL fluid in mice with or without LPS challenge. B) Mice were injected with LPS, followed by treatment with VEGF-D (225 μg/kg). The BAL fluid and histological samples were collected after 18 hours of treatment. C) The FITC-albumin probe was injected retro-orbitally 16 hours after the LPS challenge. The amount of probe in BAL fluid was quantified after another 2 hours. The FITC-albumin levels in BAL fluid in the control and experimental groups were determined. D) The BAL fluid was spun down at 5,000 g for 10 minutes and resuspended in 300 μL of 1XPBS. The number of cells in the BAL from different conditions was determined using a hemacytometer. n = 12 – 14 per group. E) LPS and LPS+VEGFD lungs were excised, fixed in 4% paraformaldehyde, embedded in paraffin, and used for histochemical analysis after haematoxylin and eosin staining. Images are representative of 6–9 lung specimens for each condition. Scale bar 150 μm. F) Lung pathological index was performed by a blinded reviewer from 1 to 4, where 4 was the worst pathology. The score was used for both peribronchial and alveolar inflammation. The average of two scores was taken to obtain a range between 1 (no inflammation or pathology) and 4 (maximum inflammation and pathology). The lung pathological index was calculated in LPS and LPS+VEGFD conditions. G-I) The expression level of TNF-α and IL-6 were measured in BAL fluid from LPS and LPS+VEGFD conditions using corresponding ELISA kits. Mice without LPS challenge were used as control. n = 12 – 14 per treatment group. Data presented as mean ± SEM. * and ** indicate p<0.05 and p<0.01, respectively.

To further explore the therapeutic potential of VEGF-D in preserving lung vascular integrity during injury, C57Bl/6 mice were randomized and subjected to intratracheal LPS injection (0.25 mg/kg) to induce inflammation. Mice were administered either a single dose of recombinant mouse VEGF-D protein or a saline control retro-orbitally following LPS instillation. VEGF-D treatment significantly reduced the levels of FITC-dextran probes in LPS-challenged animals (7339 ± 2510 (LPS) v.s. 5350 ± 1821 (LPS + VEGF-D), *p* < 0.05) (Fig. 5C).

Additionally, VEGF-D treatment resulted in a notable reduction in white blood cell count (0.791 ± 0.199 x 10^6^ WBC/mL (LPS) v.s. 0.540 ± 0.190 x 10^6^ WBC/mL (LPS + VEGF-D), *p* < 0.01) and lung injury index scores (1.518 ± 0.373 (LPS) v.s. 1.250 ± 0.250 (LPS + VEGF-D), *p* < 0.05) (Figs. 5D, E, F), underscoring VEGF-D’s role in resolving inflammation. To further characterize the impact of VEGF-D on cytokine and chemokine expression in BAL fluid, we quantified the expression levels of cytokines using ELISA. VEGF-D treatment significantly reduced the expression level of pro-inflammatory markers TNF-α and IL-6 (Figs. 5G and H) in the BAL fluid, further supporting VEGF-D’s role in mitigating inflammation. This data reveals that, consistent with findings from the in vitro cell culture model, VEGF-D protects vascular integrity and resolves increased inflammation in LPS-induced lung injury animal models.

### 3.6. VEGF-D regulates endothelial integrity through an VEGFR-2-dependent mechanism

To determine the signaling transduction on how VEGF-D regulates vascular barrier function and anti-inflammatory response, we used VEGFR2 inhibitors (SU5416, XL184, and SU5614) and siRNA transfection targeting VEGFR2 in HLMVECs. Inhibition or knockdown of VEGFR2 significantly attenuated the VEGF-D-induced enhancement of the vascular barrier (Figs. 6A and B, and Supplemental Fig. 6). The protein expression of ICAM1 and IκBα in response to VEGF-D was also diminished after VEGFR2 knockdown, indicating that VEGFR2 plays a crucial role in the anti-inflammatory effects of VEGF-D (Fig. 6C).

**Figure 6.**
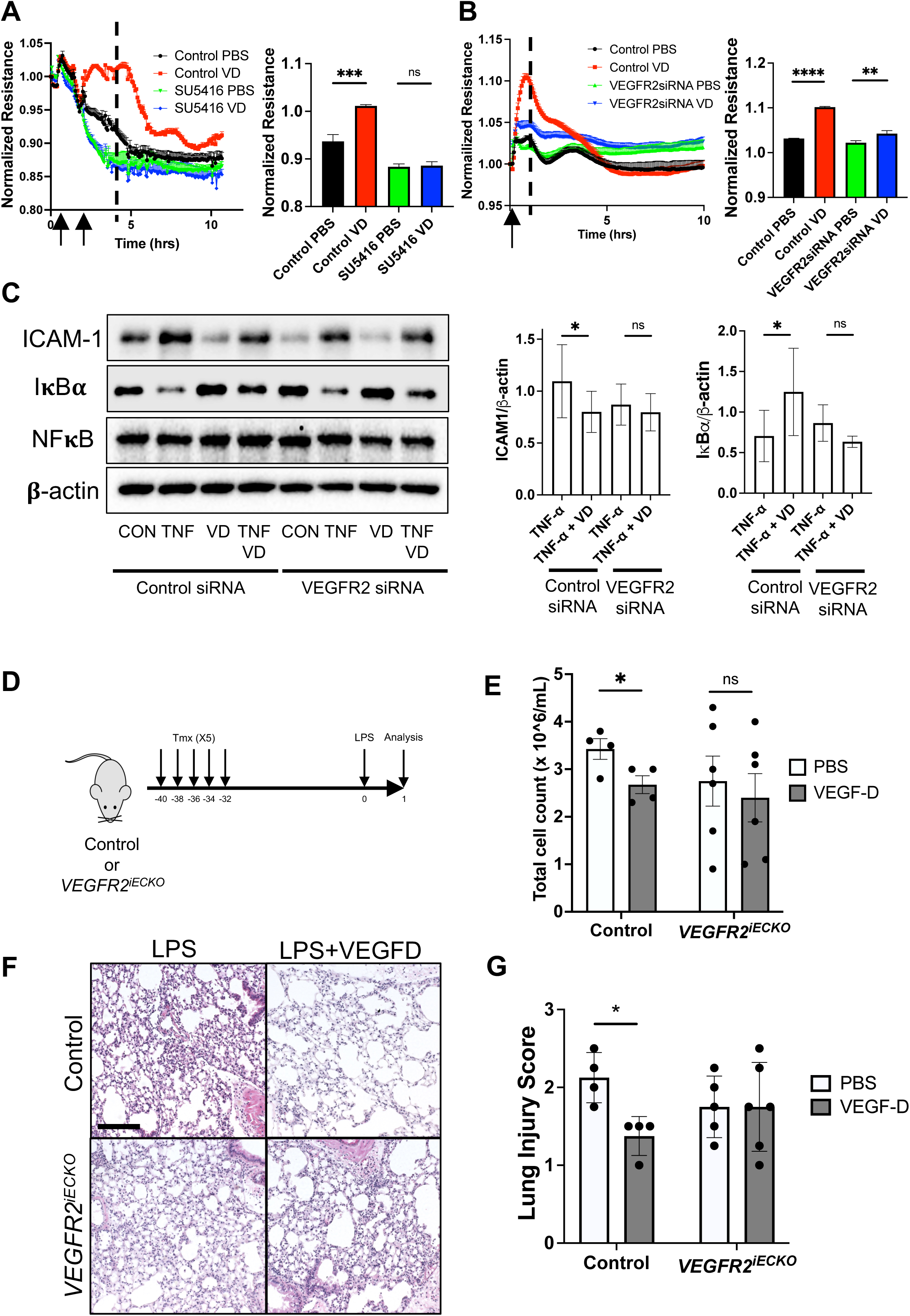
VEGF-D regulates vascular integrity through a VEGFR2-dependent mechanism. A) HLMVECs were cultured in an ECIS plate for 48 hours to reach the plateau, followed by SU5416 (5 μM) treatment. After another 1 hour, cells were treated with VEGF-D (2 μg/mL). The electrical resistance was monitored in real-time for 10 hours after drug treatment. Cells without drug treatment were used as controls. B) HLMVECs were transfected with VEGFR2 siRNA or control siRNA, then replating into ECIS plate. After reaching a plateau, cells were treated with VEGF-D or PBS. Electrical resistances were monitored. Arrows indicate drug treatment. C) HLMVECs were transfected with VEGFR2, and control siRNA followed by re-seeding into a new plate. Cells were treated with VEGF-D or PBS for 45 minutes, followed by a TNF-α (1 ng/ mL) challenge for another 5.5 hours. The protein expression of ICAM-1, IκBα, and NFκB in all conditions was determined through western blot. D-E) control (*VEGFR2^fl/fl^)* or *VEGFR2^iECKO^* at age of the 6 – 8 weeks were injected with tamoxifen 5 times on alternative days. After resting for 30 – 32 days after the last injection, mice were challenged with intratracheal injection of LPS (0.25 mg/kg) with concurrent treatment with VEGF-D (225 μg/kg). After 18 hours of injury, the total number of WBCs were counted in the BAL fluid across different experimental conditions. H) The H/E staining of lungs from control (*VEGFR2^fl/fl^)* or *VEGFR2^iECKO^* mice followed by LPS and LPS+VEGFD treatment. Images are representative of 6–9 lung specimens for each condition. Scale bar 150 μm. I) The lung pathological index was calculated in LPS and LPS +VEGFD conditions between control and *VEGFR2^iECKO^* mice. n = 4 – 6 per treatment group. Data presented as mean ± SEM. * indicates p<0.05.

To further determine the role of VEGFR2 in VEGFD signaling, we used transgenic mice with conditional deletion of VEGFR2 in endothelial cells (*VEGFR2^iECKO^*)(22, 23), a general gift from Dr. Michael Simons’ lab (49). Either control or *VEGFR2^iECKO^* mice at the age of 6 – 8 weeks were injected with tamoxifen 5 times every other day starting from 40 days prior to LPS injury. After LPS challenge, mice were injected with saline, or VEGF-D retro-orbitally and the lung tissues and BAL were harvested after another 18 hours for further analysis (Fig. 6D). In control mice, treatment with VEGF-D reduced white blood cell count in BAL and decreased lung pathological index scores after challenging with LPS, indicating reduced inflammation.

However, these anti-inflammatory effects of VEGF-D were not observed in *VEGFR2^iECKO^* mice (Figs. 6E-G) with no significant difference in WBC count, and lung pathological index score, suggesting that regulation of vascular integrity and inflammation by VEGF-D might be through a VEGFR2 dependent mechanism. These results determine the importance of VEGFR2 in mediating the effects of VEGF-D on vascular barrier function and inflammation.

## 4. Discussion

Maintaining pulmonary endothelial homeostasis is crucial for the proper functioning of the pulmonary vascular system and the preservation of the barrier integrity between the alveoli and the bloodstream. Paracrine signaling is enriched throughout the lung tissue and plays a pivotal role in regulating pulmonary vascular functions. In this study, we integrated scRNAseq analysis with biochemical validation assays to profile L/R interactions within the lung microvascular niche, specifically on datasets from healthy adult human lungs.

Focusing on the interactions between aerocyte capillary ECs and alveolar fibroblasts, we identified a putatively strong interaction between *VEGFD*, expressed by AF, and *VEGFR2* in aCap. Furthermore, we uncovered an unexpected protective role of VEGFD in maintaining vascular integrity and enhancing anti-inflammatory responses following cytokine challenges.

This suggests the VEGFD might be a potentially novel therapeutic target for treating diseases associated with lung vascular leakage, such as ALI or ARDS.

During tissue homeostasis, cell-cell crosstalk interactions within microvascular niche orchestrate vascular integrity and exert anti-inflammatory and anti-coagulation effects. Our data reveals that epithelial cells (AT1 and AT2) secrete paracrine factors that regulate neovascularization and vasodilation in microvascular endothelial cells (gCap and aCap), evidencing from significant L/R interactions, including *VEGFA-KDR*, *EDN1-EDNRB*, and *CALM1-VIPR1*. These findings align with previous observations that the pan-epithelial deletion of *Vegfa*, led to a marked decrease in aCap formation, with modest reductions in overall lung vessels and no decrease in overall endothelial cell proliferation (50).

We found that L/R interactions between mesenchymal cells (AFs, pericytes, and mesothelial cells) and microvascular endothelium primarily concentrate on signals related to vascular barrier integrity and inflammatory suppression. Notable canonical interactions include *SLIT2-ROBO4, ADM-RAMP2, and ANGPT1*-*TEK*, with particularly strong interactions between AFs and microvascular endothelium. Anatomically, these AFs have been shown to provide a bridge between AT2 cells and capillary endothelial cells, secrete paracrine signals and extracellular matrix to maintain homeostasis and facilitate remodeling during injury (51). A recent seminal study by Tsukui et al. introduced a lineage-tracing tool to fate map AF population in the lung (47). They demonstrated that ablation of AFs significantly increased IgM in BAL, neutrophil infiltration, vascular permeability and inflammatory response after bleomycin-induced lung injury (47), highlighting the protective role of AF in vascular homeostasis. Leveraging their scRNAseq data, we determined that the soluble ligands highly expressed during homeostasis including *SLIT2*, *ANGPT1*, and *VEGFD* are significantly reduced during bleomycin-induced lung injury overtime, suggesting that the protective role of AF on vasculature might be through a paracrine fashion.

In this study, we identified that VEGF-D is a key ligand produced predominantly by AFs in the distal lung of both humans and mice and interacts with microvascular ECs in the alveolar region to maintain vascular integrity. In the bleomycin-induced lung injury model, *Vegfd* expression progressively decreased as AFs transitioned from a normal to a pro-inflammatory and ultimately fibrotic phenotype. This pattern is consistent with observations from acute lung inflammation models, where *Vegfd* levels significantly declined following transient LPS treatment in both small and large animal models (52). In patients with lung diseases, such as COVID19, idiopathic pulmonary fibrosis (IPF) and systemic sclerosis-interstitial lung disease (SSc-ILD), both *VEGFD* gene expression and protein levels are markedly reduced (53–57).

Additionally, VEGF-D was shown to mediate lung fibroblast proliferation and reduce myofibroblast transition upon TGF-β stimulation (54, 58), indicating a potential role in maintaining alveolar fibroblast quiescence and contributing to the overall regulation of the alveolar niche. These findings suggest that a decrease in VEGF-D may be linked to the dysregulation of fibroblast activity and the progression of lung disease, pointing toward its broader importance in lung homeostasis and disease pathology.

We further assessed the biological impact of VEGF-D on vascular behaviors and found that VEGF-D significantly enhances lung microvascular endothelial cell barrier function and reduces the pro-inflammatory response to cytokine challenges in vitro. Additionally, we showed that VEGF-D exerted a protective effect in reducing vascular permeability and inflammation following its delivery in an LPS-induced acute lung injury model. We further demonstrated that VEGF-D’s effect on vascular integrity is through a VEGFR2-dependent mechanism in both cell culture and animal models. Notably, previous studies have suggested that VEGF-D is a potent factor in promoting angiogenesis in both lymphatic and blood vessels in skeletal muscle and heart and may induce vascular leakage in in models of hyperoxic acute lung injury (40, 41, 59). These discrepancies may be attributed to the treatment windows used in earlier research, where prolonged treatments in viral gene delivery or genetic knockout models might involve contributions from lymphatics, vessel growth, and compensatory effects from other molecules. In contrast, our study utilized the exogenous delivery of recombinant VEGF-D protein in vivo, assessing its effects within 24 hours, thereby allowing us to directly observe the impact of VEGF-D on the blood vascular barrier.

VEGFD is a secreted glycoprotein known for activating both VEGFR3 in lymphatic vessels and VEGFR2 in blood vessels (40, 41, 60). We found that VEGF-D regulates blood vascular integrity and anti-inflammatory response through a VEGFR2-dependent mechanism. Notably, the stimulation of VEGFR2 tyrosine phosphorylation by VEGFA could lead to severe vascular leakage, which is opposite to the effect of VEGFD. It was shown that VEGFD has slower kinetics in the stimulation of VEGFR2 activation and downstream signaling cascade, while the effects last longer than those induced by VEGFA (40, 41, 61, 62), which might be related to the opposite effect seen on vascular barrier regulation. However, the molecular mechanism of how VEGF-D regulates VEGFR2 to improve vascular barrier would require future extensive investigation. In conclusion, this study provides an unexpected and possible novel mechanism on the protective role of VEGFD in lung vascular integrity against injury. In combination with the previous evidence of VEGFD in alveolar fibroblast quiescence, this data sheds new light on the paracrine signal interaction in alveolar fibroblast niche, such as VEGF-D/VEGFR2, in orchestrating lung vascular homeostasis.

## 5. Acknowledgements

We thank Dr. Michael Simons and Dr. Dongying Chen at Yale University School of Medicine for providing the *VEGFR2^iECKO^* mice used in this study. We thank Dr. Yue Li for the guidance on the western blot experiments. The work on this study was supported by grants from NIH R01HL127349 (N.K.), K99/R00HL159261 (Y.Y.), UL1TR003098 (Y.Y.), AHA postdoctoral fellowships 20POST35210709 (Y.Y.), R01HL148819 (L.E.N.), an unrestricted Research Gift from Humacyte, Inc. (L.E.N.), and T32GM086287 (M.S.B.R).

## 6. Declaration of interests

In the last 3 years, N.K. served as a consultant to Biogen Idec, Boehringer Ingelheim, Third Rock, Pliant, Samumed, LifeMax, Three Lake Partners, Optikira, Astra Zeneca, RohBar, Veracyte, Augmanity, CSL Behring, Galapagos, Sofinnova, and Thyron; reports equity in Pliant and Thyron; reports grants from Veracyte, Boehringer Ingelheim, and BMS; and reports non-financial support from MiRagen and Astra Zeneca. N.K. has IP on novel biomarkers and therapeutics in IPF licensed to Biotech and is a scientific founder of Thyron. L.E.N. is a founder and shareholder in Humacyte, Inc., which is a regenerative medicine company. Humacyte produces engineered blood vessels from allogeneic smooth muscle cells for vascular surgery. L.E.N.’s spouse has equity in Humacyte, and L.E.N. serves on Humacyte’s Board of Directors. L.E.N. is an inventor on patents that are licensed to Humacyte and that produce royalties for L.E.N. L.E.N. has received an unrestricted research gift to support research in her laboratory at Yale. Humacyte did not influence the conduct, description, or interpretation of the findings in this report. The other authors report no conflicts.

## 7. Author contributions

Conceptualization, Y.Y., N.K., L.E.N., and D.W., Methodology, Y.Y., N.K., D.W., L.S., T.W., and K.G.B., Experimentation, Y.Y., L.S., T.W., Y.Y., S.K., M.S.B.R., F.A., J.K., Y.K., and Y.L., Writing – Review & Editing, Y.Y., N.K., K.G.B., D.W., M.S.B.R., Y.Y., S.K., and L.C., Discussion, Y.Y., N.K., L.E.N., D.W., K.G.B., Y.K., Q.H., Q.Y., M.S., and L.C., Funding Acquisition, Y.Y., N.K., L.E.N.

## Notes

### Competing Interest Statement

The authors have declared no competing interest.

### Summary of Updates

Acknowledgment section updated; Contribution and funding information updated

## References

1. Huertas A, Guignabert C, Barbera JA, Bartsch P, Bhattacharya J, Bhattacharya S, et al. Pulmonary vascular endothelium: the orchestra conductor in respiratory diseases: Highlights from basic research to therapy. Eur Respir J. 2018;51(4).

2. Borek I, Birnhuber A, Voelkel NF, Marsh LM, Kwapiszewska G. The vascular perspective on acute and chronic lung disease. J Clin Invest. 2023;133(16).

3. Bossardi Ramos R, Adam AP. Molecular Mechanisms of Vascular Damage During Lung Injury. Adv Exp Med Biol. 2021;1304:95–107.

4. Eilken HM, Dieguez-Hurtado R, Schmidt I, Nakayama M, Jeong HW, Arf H, et al. Pericytes regulate VEGF-induced endothelial sprouting through VEGFR1. Nat Commun. 2017;8(1):1574.

5. Rambol MH, Han E, Niklason LE. Microvessel Network Formation and Interactions with Pancreatic Islets in Three-Dimensional Chip Cultures. Tissue Eng Part A. 2020;26(9-10):556–68.

6. Doi R, Tsuchiya T, Mitsutake N, Nishimura S, Matsuu-Matsuyama M, Nakazawa Y, et al. Transplantation of bioengineered rat lungs recellularized with endothelial and adipose-derived stromal cells. Sci Rep. 2017;7(1):8447.

7. Ren X, Tapias LF, Jank BJ, Mathisen DJ, Lanuti M, Ott HC. Ex vivo non-invasive assessment of cell viability and proliferation in bio-engineered whole organ constructs. Biomaterials. 2015;52:103–12.

8. Wettschureck N, Strilic B, Offermanns S. Passing the Vascular Barrier: Endothelial Signaling Processes Controlling Extravasation. Physiol Rev. 2019;99(3):1467–525.

9. Kim H, Liu Y, Kim J, Kim Y, Klouda T, Fisch S, et al. Pericytes contribute to pulmonary vascular remodeling via HIF2alpha signaling. EMBO Rep. 2024;25(2):616–45.

10. Raredon MSB, Adams TS, Suhail Y, Schupp JC, Poli S, Neumark N, et al. Single-cell connectomic analysis of adult mammalian lungs. Sci Adv. 2019;5(12):eaaw3851.

11. Browaeys R, Saelens W, Saeys Y. NicheNet: modeling intercellular communication by linking ligands to target genes. Nat Methods. 2020;17(2):159–62.

12. Jin S, Guerrero-Juarez CF, Zhang L, Chang I, Ramos R, Kuan CH, et al. Inference and analysis of cell-cell communication using CellChat. Nat Commun. 2021;12(1):1088.

13. Schupp JC, Adams TS, Cosme C, Jr., Raredon MSB, Yuan Y, Omote N, et al. Integrated Single-Cell Atlas of Endothelial Cells of the Human Lung. Circulation. 2021;144(4):286–302.

14. Greaney AM, Adams TS, Brickman Raredon MS, Gubbins E, Schupp JC, Engler AJ, et al. Platform Effects on Regeneration by Pulmonary Basal Cells as Evaluated by Single-Cell RNA Sequencing. Cell Rep. 2020;30(12):4250–65 e6.

15. Travaglini KJ, Nabhan AN, Penland L, Sinha R, Gillich A, Sit RV, et al. A molecular cell atlas of the human lung from single-cell RNA sequencing. Nature. 2020;587(7835):619-25.

16. Guo M, Morley MP, Jiang C, Wu Y, Li G, Du Y, et al. Guided construction of single cell reference for human and mouse lung. Nat Commun. 2023;14(1):4566.

17. Raredon MSB, Yang J, Garritano J, Wang M, Kushnir D, Schupp JC, et al. Computation and visualization of cell-cell signaling topologies in single-cell systems data using Connectome. Sci Rep. 2022;12(1):4187.

18. Raredon MSB, Yang J, Kothapalli N, Lewis W, Kaminski N, Niklason LE, et al. Comprehensive visualization of cell-cell interactions in single-cell and spatial transcriptomics with NICHES. Bioinformatics. 2023;39(1).

19. Yuan Y, Leiby KL, Greaney AM, Raredon MSB, Qian H, Schupp JC, et al. A Pulmonary Vascular Model From Endothelialized Whole Organ Scaffolds. Front Bioeng Biotechnol. 2021;9:760309.

20. Yuan Y, Engler AJ, Raredon MS, Le A, Baevova P, Yoder MC, et al. Epac agonist improves barrier function in iPSC-derived endothelial colony forming cells for whole organ tissue engineering. Biomaterials. 2019;200:25–34.

21. Sharma L, Wu J, Patel V, Sitapara R, Rao NV, Kennedy TP, et al. Partially-desulfated heparin improves survival in Pseudomonas pneumonia by enhancing bacterial clearance and ameliorating lung injury. J Immunotoxicol. 2014;11(3):260–7.

22. Sun Z, Li X, Massena S, Kutschera S, Padhan N, Gualandi L, et al. VEGFR2 induces c-Src signaling and vascular permeability in vivo via the adaptor protein TSAd. J Exp Med. 2012;209(7):1363–77.

23. Wang Y, Nakayama M, Pitulescu ME, Schmidt TS, Bochenek ML, Sakakibara A, et al. Ephrin-B2 controls VEGF-induced angiogenesis and lymphangiogenesis. Nature. 2010;465(7297):483-6.

24. Braud VM, Allan DS, O’Callaghan CA, Soderstrom K, D’Andrea A, Ogg GS, et al. HLA-E binds to natural killer cell receptors CD94/NKG2A, B and C. Nature. 1998;391(6669):795–9.

25. Dostert C, Grusdat M, Letellier E, Brenner D. The TNF Family of Ligands and Receptors: Communication Modules in the Immune System and Beyond. Physiol Rev. 2019;99(1):115–60.

26. Gillich A, Zhang F, Farmer CG, Travaglini KJ, Tan SY, Gu M, et al. Capillary cell-type specialization in the alveolus. Nature. 2020;586(7831):785–9.

27. London NR, Zhu W, Bozza FA, Smith MC, Greif DM, Sorensen LK, et al. Targeting Robo4-dependent Slit signaling to survive the cytokine storm in sepsis and influenza. Sci Transl Med. 2010;2(23):23ra19.

28. Mei SH, McCarter SD, Deng Y, Parker CH, Liles WC, Stewart DJ. Prevention of LPS-induced acute lung injury in mice by mesenchymal stem cells overexpressing angiopoietin 1. PLoS Med. 2007;4(9):e269.

29. Shrestha AK, Menon RT, Yallampalli C, Barrios R, Shivanna B. Adrenomedullin Deficiency Potentiates Lipopolysaccharide-Induced Experimental Bronchopulmonary Dysplasia in Neonatal Mice. Am J Pathol. 2021;191(12):2080–90.

30. Menon RT, Shrestha AK, Reynolds CL, Barrios R, Caron KM, Shivanna B. Adrenomedullin Is Necessary to Resolve Hyperoxia-Induced Experimental Bronchopulmonary Dysplasia and Pulmonary Hypertension in Mice. Am J Pathol. 2020;190(3):711–22.

31. Herradon G, Perez-Garcia C. Targeting midkine and pleiotrophin signalling pathways in addiction and neurodegenerative disorders: recent progress and perspectives. Br J Pharmacol. 2014;171(4):837–48.

32. Lazarus A, Keshet E. Vascular endothelial growth factor and vascular homeostasis. Proc Am Thorac Soc. 2011;8(6):508–11.

33. Markovic R, Peltan J, Gosak M, Horvat D, Zalik B, Seguy B, et al. Planar cell polarity genes frizzled4 and frizzled6 exert patterning influence on arterial vessel morphogenesis. PLoS One. 2017;12(3):e0171033.

34. Rickman M, Ghim M, Pang K, von Huelsen Rocha AC, Drudi EM, Sureda-Vives M, et al. Disturbed flow increases endothelial inflammation and permeability via a Frizzled-4-beta-catenin-dependent pathway. J Cell Sci. 2023;136(6).

35. Nabhan AN, Webster JD, Adams JJ, Blazer L, Everrett C, Eidenschenk C, et al. Targeted alveolar regeneration with Frizzled-specific agonists. Cell. 2023;186(14):2995–3012 e15.

36. Broderick TL, Wang Y, Gutkowska J, Wang D, Jankowski M. Downregulation of oxytocin receptors in right ventricle of rats with monocrotaline-induced pulmonary hypertension. Acta Physiol (Oxf). 2010;200(2):147–58.

37. Weng J, Zhou X, Xie H, Gao Y, Wang Z, Gong Y. Slit2/Robo4 signaling pathway modulates endothelial hyper-permeability in a two-event in vitro model of transfusion-related acute lung injury. Blood Cells Mol Dis. 2019;76:7–12.

38. Ismail H, Mofarrahi M, Echavarria R, Harel S, Verdin E, Lim HW, et al. Angiopoietin-1 and vascular endothelial growth factor regulation of leukocyte adhesion to endothelial cells: role of nuclear receptor-77. Arterioscler Thromb Vasc Biol. 2012;32(7):1707–16.

39. Geven C, Kox M, Pickkers P. Adrenomedullin and Adrenomedullin-Targeted Therapy As Treatment Strategies Relevant for Sepsis. Front Immunol. 2018;9:292.

40. Rutanen J, Leppanen P, Tuomisto TT, Rissanen TT, Hiltunen MO, Vajanto I, et al. Vascular endothelial growth factor-D expression in human atherosclerotic lesions. Cardiovasc Res. 2003;59(4):971–9.

41. Rutanen J, Rissanen TT, Markkanen JE, Gruchala M, Silvennoinen P, Kivela A, et al. Adenoviral catheter-mediated intramyocardial gene transfer using the mature form of vascular endothelial growth factor-D induces transmural angiogenesis in porcine heart. Circulation. 2004;109(8):1029–35.

42. Prasain N, Stevens T. The actin cytoskeleton in endothelial cell phenotypes. Microvasc Res. 2009;77(1):53–63.

43. Shen Q, Rigor RR, Pivetti CD, Wu MH, Yuan SY. Myosin light chain kinase in microvascular endothelial barrier function. Cardiovasc Res. 2010;87(2):272–80.

44. Collins T, Read MA, Neish AS, Whitley MZ, Thanos D, Maniatis T. Transcriptional regulation of endothelial cell adhesion molecules: NF-kappa B and cytokine-inducible enhancers. FASEB J. 1995;9(10):899–909.

45. Bastarache JA, Blackwell TS. Development of animal models for the acute respiratory distress syndrome. Dis Model Mech. 2009;2(5-6):218–23.

46. Ghonim MA, Boyd DF, Flerlage T, Thomas PG. Pulmonary inflammation and fibroblast immunoregulation: from bench to bedside. J Clin Invest. 2023;133(17).

47. Tsukui T, Wolters PJ, Sheppard D. Alveolar fibroblast lineage orchestrates lung inflammation and fibrosis. Nature. 2024;631(8021):627–34.

48. Tsukui T, Sun KH, Wetter JB, Wilson-Kanamori JR, Hazelwood LA, Henderson NC, et al. Collagen-producing lung cell atlas identifies multiple subsets with distinct localization and relevance to fibrosis. Nat Commun. 2020;11(1):1920.

49. Chen D, Rukhlenko OS, Coon BG, Joshi D, Chakraborty R, Martin KA, et al. VEGF counteracts shear stress-determined arterial fate specification during capillary remodeling. bioRxiv. 2024.

50. Vila Ellis L, Cain MP, Hutchison V, Flodby P, Crandall ED, Borok Z, et al. Epithelial Vegfa Specifies a Distinct Endothelial Population in the Mouse Lung. Dev Cell. 2020;52(5):617–30 e6.

51. Sirianni FE, Chu FS, Walker DC. Human alveolar wall fibroblasts directly link epithelial type 2 cells to capillary endothelium. Am J Respir Crit Care Med. 2003;168(12):1532–7.

52. Toth A, Steinmeyer S, Kannan P, Gray J, Jackson CM, Mukherjee S, et al. Inflammatory blockade prevents injury to the developing pulmonary gas exchange surface in preterm primates. Sci Transl Med. 2022;14(638):eabl8574.

53. Adams TS, Schupp JC, Poli S, Ayaub EA, Neumark N, Ahangari F, et al. Single-cell RNA-seq reveals ectopic and aberrant lung-resident cell populations in idiopathic pulmonary fibrosis. Sci Adv. 2020;6(28):eaba1983.

54. Cui Y, Osorio JC, Risquez C, Wang H, Shi Y, Gochuico BR, et al. Transforming growth factor-beta1 downregulates vascular endothelial growth factor-D expression in human lung fibroblasts via the Jun NH2-terminal kinase signaling pathway. Mol Med. 2014;20(1):120–34.

55. Reyfman PA, Walter JM, Joshi N, Anekalla KR, McQuattie-Pimentel AC, Chiu S, et al. Single-Cell Transcriptomic Analysis of Human Lung Provides Insights into the Pathobiology of Pulmonary Fibrosis. Am J Respir Crit Care Med. 2019;199(12):1517–36.

56. Renaud L, Wilson CL, Lafyatis R, Schnapp LM, Feghali-Bostwick CA. Transcriptomic characterization of lung pericytes in systemic sclerosis-associated pulmonary fibrosis. iScience. 2024;27(6):110010.

57. Sikkema L, Ramirez-Suastegui C, Strobl DC, Gillett TE, Zappia L, Madissoon E, et al. An integrated cell atlas of the lung in health and disease. Nat Med. 2023;29(6):1563–77.

58. El-Chemaly S, Pacheco-Rodriguez G, Malide D, Meza-Carmen V, Kato J, Cui Y, et al. Nuclear localization of vascular endothelial growth factor-D and regulation of c-Myc-dependent transcripts in human lung fibroblasts. Am J Respir Cell Mol Biol. 2014;51(1):34–42.

59. Sato T, Paquet-Fifield S, Harris NC, Roufail S, Turner DJ, Yuan Y, et al. VEGF-D promotes pulmonary oedema in hyperoxic acute lung injury. J Pathol. 2016;239(2):152–61.

60. Rissanen TT, Markkanen JE, Gruchala M, Heikura T, Puranen A, Kettunen MI, et al. VEGF-D is the strongest angiogenic and lymphangiogenic effector among VEGFs delivered into skeletal muscle via adenoviruses. Circ Res. 2003;92(10):1098–106.

61. Toivanen PI, Nieminen T, Viitanen L, Alitalo A, Roschier M, Jauhiainen S, et al. Novel vascular endothelial growth factor D variants with increased biological activity. J Biol Chem. 2009;284(23):16037–48.

62. Jia YT, Li ZX, He YT, Liang W, Yang HC, Ma HJ. Expression of vascular endothelial growth factor-C and the relationship between lymphangiogenesis and lymphatic metastasis in colorectal cancer. World J Gastroenterol. 2004;10(22):3261–3.

