## SupplementalFigures for "Vascular endothelial growth factor-D improves lung vascular integrity during acute lung injury"

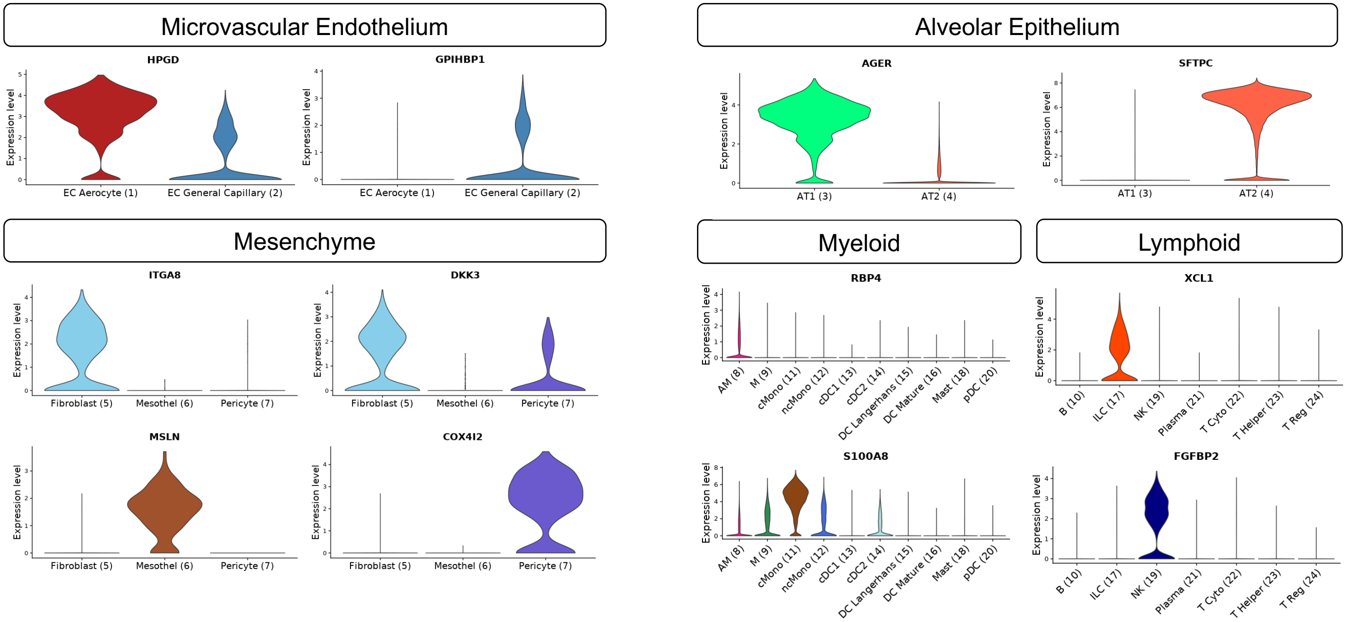


**Supplemental Figure 1: Representative markers of each cell cluster are shown in Figure 2A**. VlnPlot of representative markers for EC aerocyte (*HPGD*), EC General Capillary (*GPIHBP1*), Alveolar fibroblast (*ITGA8* and *DKK3*), Mesothelium (*MSLN*), Pericyte (*COX4I2*), AT1 (*AGER*), AT2 (*SFTPC*), AM (*RBP4*), cMono (*S100A8*), ILC (*XCL1*) and NK (*FGFBP2*) to confirm clustering and annotation of cell populations in human lung microvascular niche.

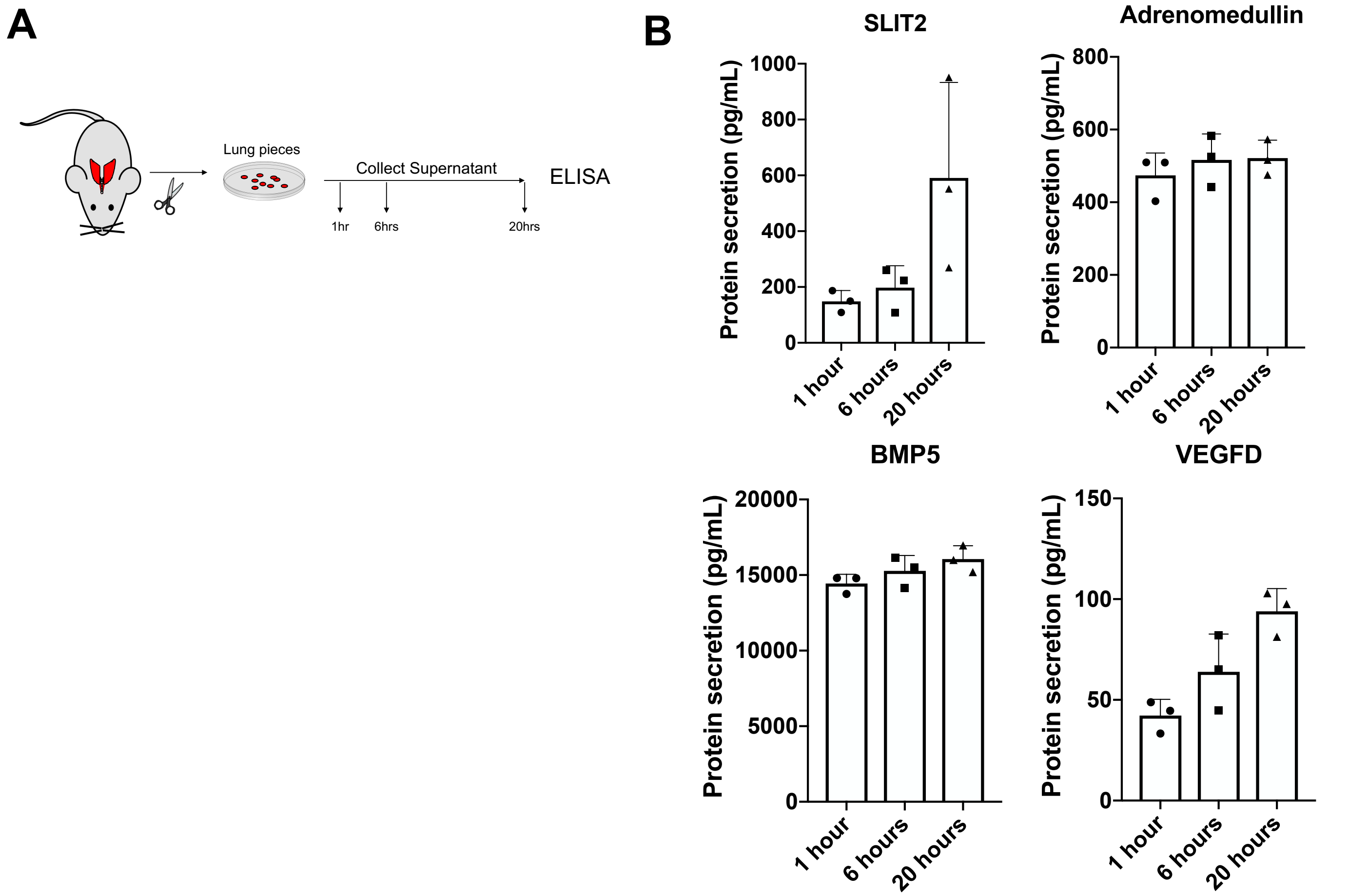


**Supplemental Figure 2: The secretion levels of identified ligands in ex vivo rat lung culture.** A) Adult rat lungs were cut into small pieces (<1 $mm^3$ per piece) and cultured into the serum-reduced culture medium for 20 hours. The supernatant of the ex vivo lung culture was collected at 1 hour, 6 hours, and 20 hours. B) Slit2, BMP5, Adrenomedullin, and VEGFD ELISA were performed using the corresponding rat ELISA kit as per manufacturer's instructions.

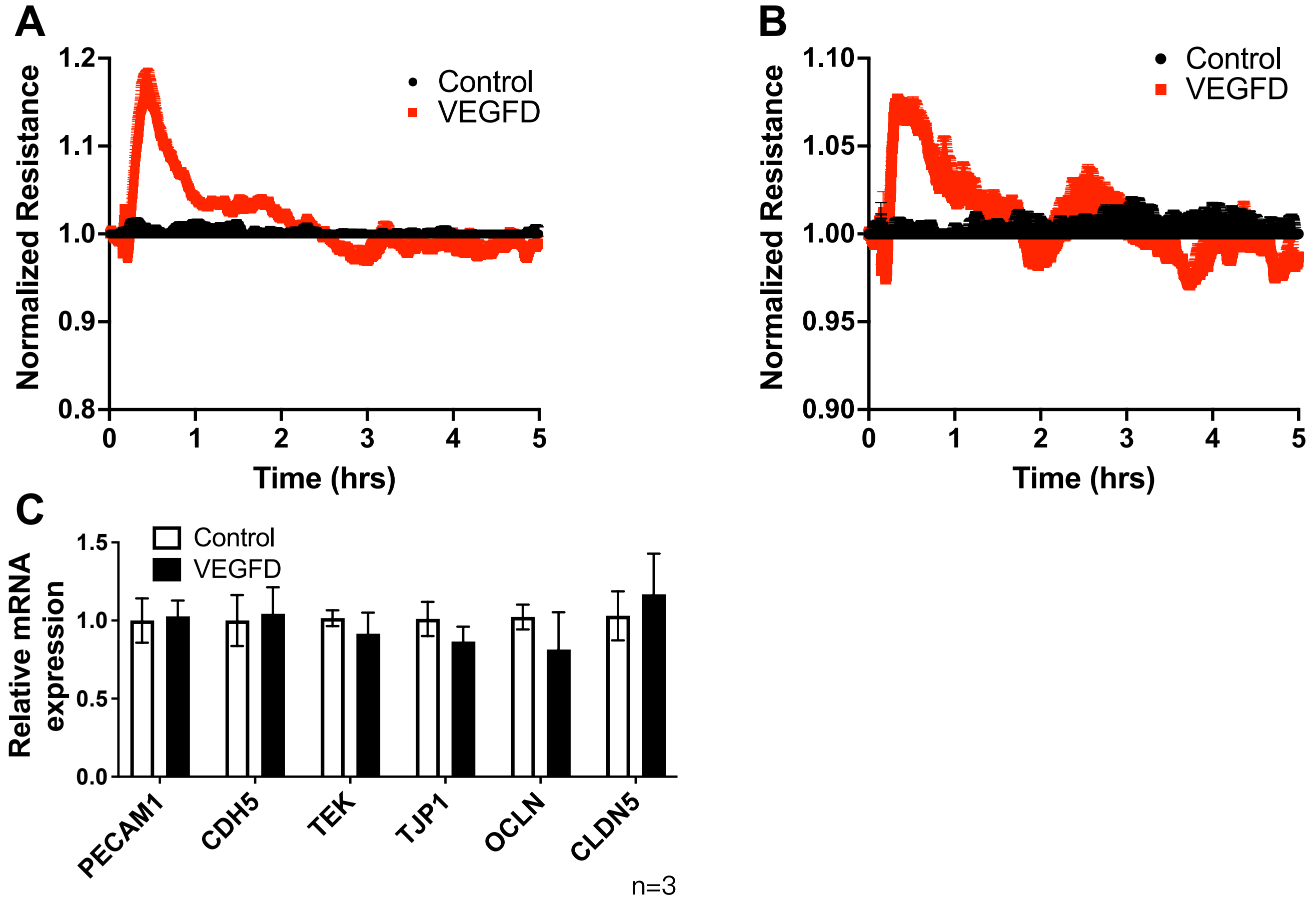


**Supplemental Figure 3: VEGF-D and barrier function in endothelial cells.** A, B) The electrical resistance in human pulmonary arterial endothelial cells (HPAECs) and human pulmonary venous endothelial cells (HPVECs) after treatment with VEGF-D (1 µg/mL) were measured through electrical cell impendence sensing (ECIS) assay. C) The expression levels of cell-cell junctional proteins *CDH5*, *TJP1*, *OCLN*, *CLDN5*, *TEK*, and *PECAM1* were measured through qPCR in HLMVECs with or without VEGF-D treatment. There were 3 replicates performed in each sample.

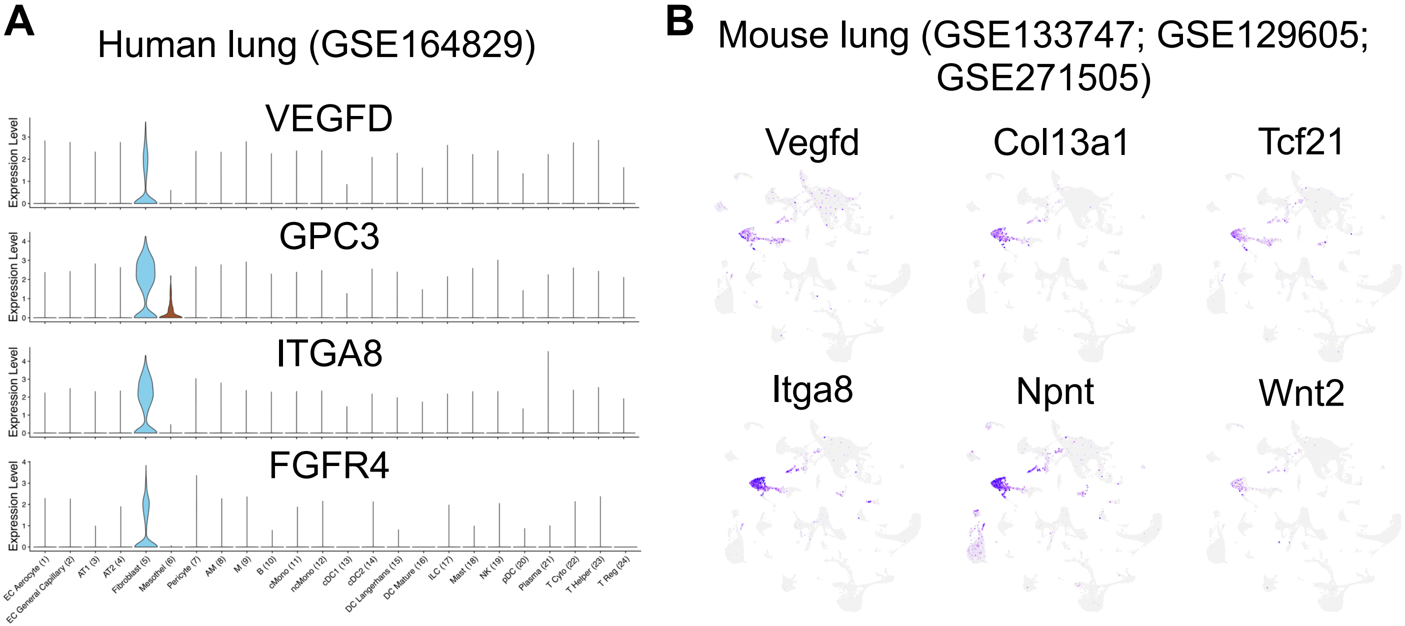


**Supplemental Figure 4: VEGF-D expresses in human and mouse alveolar fibroblast.** A) VlnPlot of *VEGFD*, *GPC3*, *ITGA8*, and *FGFR4* in human lung scRNAseq atlas (GSE164829). B) FeaturePlot of *Vegfd*, *Col13a1*, *Tcf21*, *Itga8*, *Npnt*, and *Wnt2* from an integrated scRNAseq object contains all 28935 mouse cells from 8 control mouse lungs from 3 independent studies (1-3).

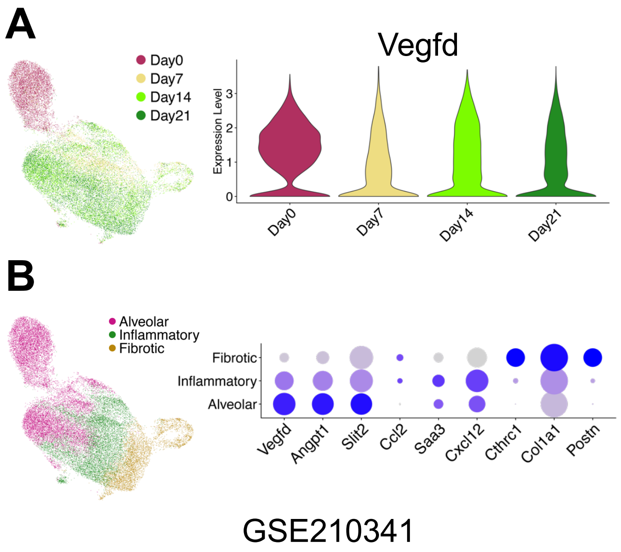


**Supplemental Figure 5: VEGF-D expression level in bleomycin-induced animal model.** A) Uniform manifold approximation and projection of a total of 28935 cells in the re-clustered tdtamado+ alveolar fibroblasts (AF) scRNAseq data in *Scube2-creER Rosa26-tdTomato* mice (GSE210341)(4). The metadata was copied from an original dataset containing information on the day's post-bleomycin treatment. VlnPlot shows the *Vegfd* expression level in the AF population on day 0, day 7, day 14, and day 21 after the bleomycin challenge. B) Uniform manifold approximation and projection of scRNAseq data of tdtamado+ alveolar fibroblasts (AF) subsetted into alveolar, inflammatory and fibrotic fibroblasts. DotPlot shows the expression level of *Vegfd*, *Angpt1*, *Slit2*, *Ccl2*, *Saa3*, *Cxcl12*, *Cthrc1*, *Col1a1*, and *Postn* in alveolar, inflammatory and fibrotic fibroblasts.

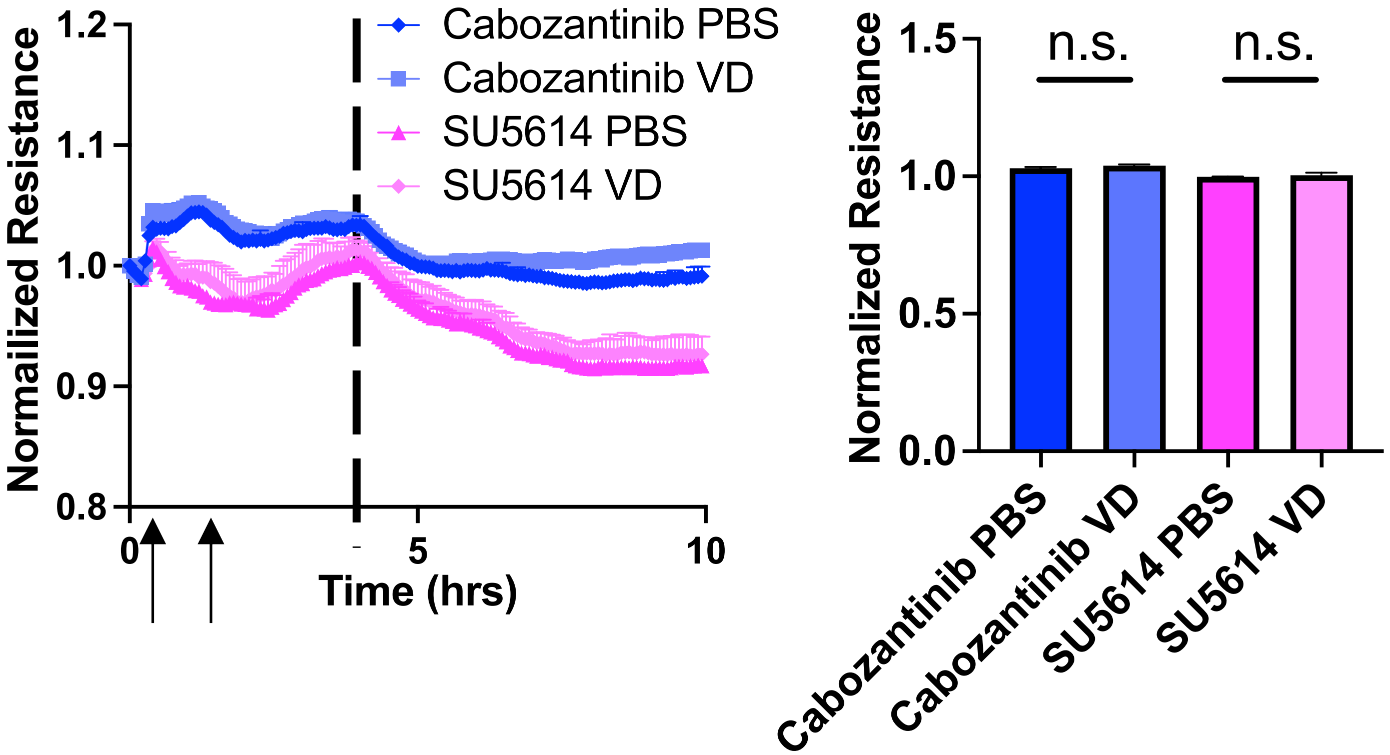


**Supplemental Figure 6: Additional VEGFR2 inhibitors and endothelial barrier function.** HLMVECs were cultured in an ECIS plate for 48 hours to reach the plateau, followed by treatment with VEGFR2 selective pharmacological inhibitors Cabozantinib (2 nM) or SU5614 (10 µM). After another 1 hour, cells were then treated with VEGF-D (2 µg/mL). The electrical resistance was monitored for the following 10 hours. Cells without drug treatment were used as controls. There are 3 – 4 experiments performed.
